# Alteration of coastal productivity and artisanal fisheries interact to affect a marine food-web

**DOI:** 10.1101/2020.10.18.343756

**Authors:** M. Isidora Ávila-Thieme, Derek Corcoran, Alejandro Pérez-Matus, Evie A. Wieters, Sergio A. Navarrete, Pablo A. Marquet, Fernanda S. Valdovinos

## Abstract

Top-down and bottom-up forces determine ecosystem function and dynamics. Fisheries as a top-down force can shorten and destabilize food-webs, while climate-change driven effects can alter the bottom-up forces of primary productivity. We assessed the response of a highly-resolved intertidal food-web to these two global-change drivers, using network analysis and bioenergetic modelling. We quantified the relative importance of artisanal fisheries as another predator species, and evaluated the independent and combined effects of fisheries and plankton-productivity changes on food-web dynamics. The food-web was robust to the loss of all harvested species but sensible to plankton-productivity decline. Interestingly, fisheries dampened the negative impacts of decreasing plankton productivity on non-harvested species, while plankton-productivity decline increased the sensitivity of harvested species to fishing. Our results show that strategies for new scenarios caused by climate change are needed to protect marine ecosystems and the wellbeing of local communities dependent on their resources.

## INTRODUCTION

Direct human impacts and the full suite of drivers of global change are the main cause of species extinctions in Anthropocene ecosystems^1,2^, with detrimental consequences on ecosystem functioning and their services to human societies^3,4^. The world fisheries crisis is among those consequences, which cuts across fishing strategies, oceanic regions, species, and includes countries that have little regulation and those that have implemented rights-based co-management strategies to reduce overharvesting^5–8^. Chile has been one of the countries implementing Territorial Use Rights (TURFs^9^) over an unprecedented geographic scale to manage the diverse coastal benthic resources using a co-management strategy^10,11^. Over 60 coastal benthic species form part of these artisanal fisheries^10^, with species that are extracted from intertidal and shallow subtidal habitats ^12,13^. The Chilean TURFs system brought significant improvements in sustainability of this complex socio-ecological system, helping to rebuild benthic fish stocks^10,11^, improving fishers’ perception towards sustainability and increasing compliance^9^, as well as showing positive ancillary effects on conservation of biodiversity^14,15^. However, the situation of most artisanal fisheries is still far from sustainable, and many fish stocks and coastal ecosystems show signs of over exploitation and ecosystem degradation, a consequence of the low levels of cooperation and low enforcement of TURF regulations, which leads to high levels of free-riding and illegal fishing^16–18^. Thus, it is imperative to improve our understanding of the effects of these multi-species fisheries which simultaneously harvest species at all trophic levels, from kelp primary producers to top carnivores^13,19^.

To compound things, removal of biomass from the ocean occurs simultaneously with multiple other stressors associated to climate change that compromise the individuals’ capacity to respond to perturbations^20–22^. Besides sea surface temperature (SST), climate change also affects many other physical-chemical characteristics of marine coastal waters (stratification, acidification, ventilation)^23,24^, as well as the wind regimes that control surface water productivity along the productive coastal upwelling ecosystems^25–29^. Changes in the productivity of the oceans are reflected in changes of plankton biomass, which contributes approximately half of the global primary production, supports the productivity of marine food-webs, and influences the biogeochemical process in the ocean and strongly affects commercial fisheries^30–32^. Indeed, an overall decrease in marine plankton productivity is expected over global scales^24,30,33^. Along extensive regions of the Humboldt upwelling ecosystem off Chile, long-term increases and decreases in plankton productivity have already occurred over the past two decades^34,35^ and are expected to propagate up the pelagic and benthic food webs. We therefore analyzed the bottom-up impact of fluctuations in plankton productivity in combination with fisheries exploitation of these food-webs using the concepts and methods of network ecology.

Network ecology has advanced our understanding of ecosystems by providing a powerful framework to analyze biological communities^36,37^. Previous studies used this framework to assess food-web robustness against species extinctions (i.e., the fraction of initial species that remain present in the ecosystem after a primary extinction) ^38–43^, showing the importance for food-web persistence of highly connected species^38,40,44,45^, basal species^39^ and highly connected species that trophically support other highly connected species^42^. Most of these studies used a static approach, which stems from network theory and analyzes the impacts of structural changes on food-webs represented by nodes (species) and links (interactions) that connect nodes, but ignores interaction strengths and population dynamics of interacting species^38^. Other studies used a dynamic approach, which considers not only the structure and intensity of interactions in a food-web, but also the changes in species biomasses through time and the indirect effects that these changes have on other species^39–41,46,47^. Here we use both approaches to understand the relative importance of harvested species in our food-web.

In this contribution, we analyze (1) the importance of harvested species for the structure and persistence of the intertidal food-web by simulating a scenario of over-exploitation-driven extinction of all harvested species. We then evaluate (2) the robustness of this food-web to the extinction of species harvested by artisanal fisheries in comparison to three commonly used extinctions sequences (see below), and (3) the effect of three fisheries scenarios on other species abundance, persistence and food-web dynamics. We finally analyze the (4) independent and (5) combined effects of fisheries and plankton productivity changes on the food-web dynamics through altering plankton subsidy.

## RESULTS

### 1. Food-web description and the relative importance of harvested species to the food-web structure

The intertidal food-web contains 107 species, with the highly omnivorous fisheries node (F node in Fig. 1A) contributing 22 links, from basal kelp species to top carnivores (Fig. 1A). Among the species harvested by the artisanal fisheries, 10 belong to the 30 most connected species of the food-web (Fig. 1A, and Supplementary Table S1). Moreover, these fisheries exploit at least one species that is a resource or a consumer of about 70% of the species (harvested and non-harvested species) in the intertidal food-web (Fig. 1B, C and D). With the static approach we found that the removal of all 22 species (see Methods) negatively affected the structural properties of the food-web, specially, reduces the overall number of trophic interactions by 48%. This loss represents, on average, 100 more links lost than that expected from randomly removing 22 species from the food-web (see supplementary Table S2 and supplementary material for more detailed results).

**Figure 1.**
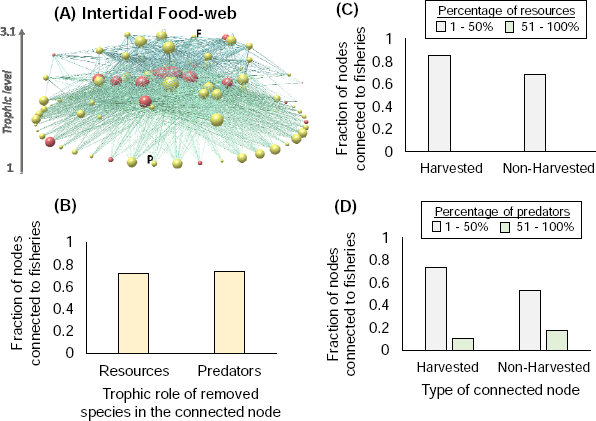
(A) Intertidal food-web. Node colors represent harvested (red) and non-harvested (yellow) species. Letter F and P represent the fisheries and plankton node, respectively. Node size represents the number of trophic interactions (degree) of each node. Nodes at the bottom represent basal species, while nodes at the top represent top predators. Y-axis represents trophic level (calculated as SWTL, see Methods). Drawn using Network3D software^48^. (B) Fraction of species in the food web that are trophically connected (at least once) with exploited species that are either a resource and/or a predator (x-axis). Each bar presented in (B) is further divided in: (C) the percentage of resources shared with fisheries by the harvested and non-harvested consumers of the food-web, and (D) the percentage of predators of harvested and non-harvested species extracted by fisheries. Grey and green bars, respectively, represent the categories of 1-50% and 51-100% of the resource species consumed by harvested and non-harvested species (C) and of the consumer species predating upon harvested and non-harvested species (D).

### 2. Food-web robustness to species extinctions

Following previous work^38–41^, we evaluated the robustness of the intertidal food-web to species extinction by sequentially removing species and counting the subsequent secondary extinctions, if any. We counted the secondary extinctions caused by the four deletion sequences (harvesting, random, most-connected, supporting-basal) using both static and dynamic approaches (see Methods). Our dynamic approach uses and extends the Allometric Trophic Network^49,50^ (ATN) model by including plankton subsidy. Both approaches found that the intertidal food web is highly robust to the loss of all harvested species, as null secondary extinctions were observed after removing all harvested species (Fig. 2). The robustness of the intertidal food-web was further demonstrated by the sequential deletion of the most connected species, which showed that over 30% of those species must be removed before any secondary extinctions occur (Fig. 2). As expected from previous work, the loss of supporting-basal species produced the most secondary extinctions (Fig. 2). Both approaches showed similar trends (Fig. 2A and B), but our dynamic approach presented relatively lower food-web robustness (Supplementary Fig. S1).

**Figure 2.**
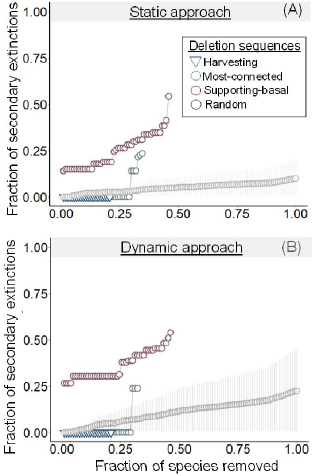
Fraction of secondary extinctions (y-axis) produced by the sequential removal of species (x-axis) with static (A) and dynamic (B) approaches. Gray and red circles represent most-connected and supporting-basal deletion sequences, while blue tringle represents harvesting deletion sequence. In the random deletion sequence, circles represent the average and the error bars represent the 95% confidence interval over 1000 simulations.

Although the plankton node (“species”) was directly connected only to filter-feeders, it proved to be the most important in the supporting-basal deletion sequence for both static and dynamic approaches, as its removal caused 15 and 29 secondary extinctions, respectively. The species that went extinct included not only sessile filter-feeders, but also four harvested species important for the fisheries: the Chilean muricid whelk *Concholepas concholepas*, the giant barnacle *Austromegabalanus pssitacus*, the sea squirt *Pyura chilensis* and the whelk *Acanthina monodon*. These results suggest that while the intertidal food-web is robust to harvest-driven extinctions, it can be sensitive to a drastic decrease in plankton productivity.

### 3. Effects of artisanal fisheries on food-web dynamics

We assessed the effects of fisheries on the biomass of every species in the food-web using our extension of the ATN model (see Methods). Fig. 3A and 3B illustrates with a simplified diagram our results shown in Supplementary Fig. S2. We simulated three fishing scenarios, where we applied exploitation rates needed to decrease the biomass of all 22 harvested species in 50%, 80%, and 100% (see F_max_ in Supplementary Table S3). These three fishing scenarios allowed us to simulate an approximately well managed fisheries (which removes between 40% and 60% of biomass stock^5^), an overexploitation scenario (which removes 80%) and nearly extinction scenario, respectively. We found that basal species required much lower exploitation rate to decrease their biomass than filter-feeders, herbivores, and other consumers (Supplementary Table S3). Harvested basals went extinct with an extraction above 0.3% of their available biomass, while harvested consumers went extinct with an extraction above 90% of their available biomass.

**Figure 3.**
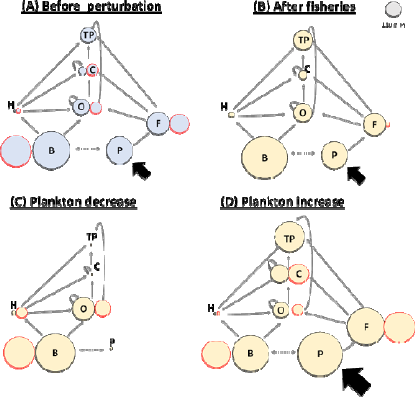
Diagram illustrating the effect of fisheries (B) and the effect of perturbing plankton subsidy (C and D) on food-web dynamics. Nodes represent the total biomass of each trophic level before (A) and after reducing in 100% the biomass of all harvested species (B) and after decreasing (C) or increasing (D) plankton subsidy in 100%. Each trophic level is indicated by TP: top predators, C: carnivores, O: omnivores, H: herbivores, F: filters-feeders, B: basal species, and P: plankton. Red and black outlined nodes represent the biomass of harvested and non-harvested species, respectively. Solid black, solid grey, and dashed grey arrows represent the plankton subsidy (7355 g/m^2^), the trophic interactions, and competitive interactions, respectively. Note that the ATN model explicitly models competition only between basal species, while competition between consumers emerges from the depletion of shared resources.

The decrease in biomass of harvested species led to an increase in the biomass of most non-harvested species at all trophic levels, especially in basal and herbivorous species (compare Figs. 3A and 3B). On average, 82-86% of non-harvested species increased their biomass by 5-25% after 50-100% fishing impacts on biomass stocks (Supplementary Fig. S2). This biomass increase is explained by two mechanisms: i) decreasing the biomass of harvested species that are consumers reduces the predation intensity on their resources (note that fisheries harvest more species in higher than lower trophic levels, compare Figs. 1D and 1C), and (ii) decreasing the biomass of harvested basal species reduces their competitive effects on the non-harvested basal species, allowing them to grow (Supplementary Fig. S3).

The positive effect of artisanal fisheries on the biomass of non-harvested species was qualitatively similar across the different fishing scenarios, becoming larger with an increase in fishing intensity (Supplementary Fig. S2). The exceptions were top predators, which had opposite responses between the weakest and strongest fishing scenarios. A 50% reduction in harvested species biomass caused a slight decrease in non-harvested top predators’ biomass, while 80% and 100% reductions caused an increase in their biomass. This suggests that artisanal fisheries negatively impact top predators by extracting their prey species but, when the exploitation rates are stronger, the indirect positive effects of fisheries on the biomass of the non-harvested species become strong enough to dampen those effects.

### 4. Effects of plankton-subsidy alteration on food-web dynamics

We considered a subsidy term to plankton as an external-controlled subsidy of plankton productivity. We both decreased (Fig. 3C) and increased (Fig. 3D) the plankton subsidy in 50%, 80%, and 100% to simulate the alteration of plankton productivity expected as a response of climate change (see Methods). All biomass changes can be found in Supplementary Fig. S4. Both decreasing (Fig. 3C) and increasing (Fig. 3D) plankton subsidy can deeply alter food web dynamics. The magnitude of plankton subsidy increases or decreases (i.e., 50%, 80%, 100%) only quantitatively affected the food-web patterns shown in Figs. 3C and 3D, becoming more intense with an increasing alteration of the plankton subsidy. Decreasing plankton subsidy had larger impacts on the species biomasses than increasing plankton subsidy in the same magnitude, even causing species extinctions (i.e., -1 in Supplementary Fig. S4E) when the subsidy was removed (i.e., decreased in 100%). The number of total extinctions that occurred after completely removing the plankton subsidy was 29 species, highlighting the bottom-up propagation of effects through the web (Supplementary Fig. S4E).

A drastic decrease in plankton subsidy (100%) resulted in the extinction of all filter-feeder species (specialist consumers of plankton) and decreased the biomass of carnivores and top predators by 99% (compare Figs. 3A and 3C). The biomass reduction in carnivores and top predators, in turn, released predation pressure on omnivores and herbivores, which consequently increased their biomass by 30% and 110%, respectively. The increased biomass of herbivores and omnivores, in turn, increased consumption pressure on basal species, but we found that the biomass of basal species slightly increased by 4% (Fig. 3C). This suggests that the reduction in plankton subsidy positively affects basal species by releasing pressure on the community level carrying capacity (see Methods). Conversely, a 100% increase in plankton subsidy increased the total biomass of filters, carnivores, and top predators by 76%, 107%, and 105%, respectively (compare Figs. 3A and 3D). As a consequence, the increased predation pressure from higher trophic levels decreased the total biomass of omnivores, herbivores, and basal species by 2%, 20%, and 3%, respectively. Carnivore species were the most vulnerable to the reduction of plankton productivity, going extinct with a reduction of 80% in plankton subsidy (Supplementary Fig. S4C), followed by filter-feeders and top predators, which went extinct with a 100% of subsidy reduction (Supplementary Fig. S4E). Regarding harvested species, 18% of them strongly decreased their biomass when plankton subsidy decreased, while 81% of them slightly decreased their biomass when plankton subsidy increased (compare Supplementary Figs. S4A, S4C, S4E with S4B, S4D, S4F).

### 5. Interacting effects of fisheries and plankton-subsidy alteration on food-web dynamics

We evaluated the combined effects of the biomass extraction by fisheries and the alteration of plankton subsidy on the food-web dynamics using a full factorial design that maintains the same fishing and subsidy levels used in each of the last two sections. We found that regardless of the fishing scenario, all non-harvested trophic levels persisted when the plankton subsidy increased or decreased (Fig. 4A and B) by 50%. Conversely, when the plankton subsidy decreased in 80%, carnivores went extinct under all fishing scenarios (compare Supplementary Figs. S5C and D with S5A, B, E and F) as well as the top-predators and filter-feeders when the plankton subsidy decreased in 100% (Fig. 4C and D).

**Figure 4.**
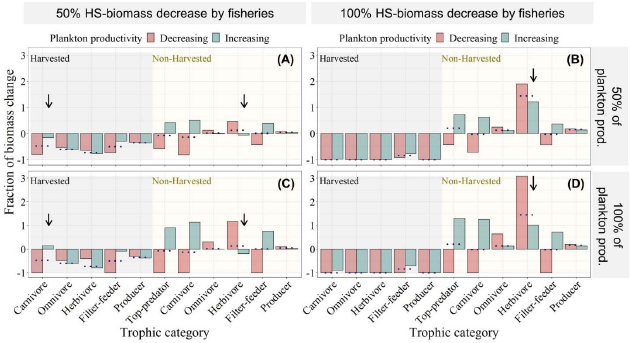
Combined effects of artisanal fisheries and plankton-productivity alterations on food-web dynamics. Fraction of total biomass change (y-axis) of each trophic category (x-axis) after decreasing (red bars) and increasing (blue bars) the plankton productivity (plankton prod.) in 50% (A and B) and 100% (C and D), and after decreasing the biomass of all harvested species (HS) in a 50% (A and C) and in a 100% (B and D). The grey and yellow shading represent the biomass change of harvested and non-harvested species, respectively. The arrows highlight the most remarkable changes between the two levels of plankton subsidy perturbation and the two levels of fishing. The dotted lines represent the independent effect fishing (i.e., without plankton subsidy perturbation) on the biomass of each trophic category as a reference point.

The level of plankton subsidy affected the impact of fishing on the biomass of harvested species. Decreasing plankton subsidy decreased the biomass of harvested carnivores and filter-feeders, intensifying the negative effect of fishing on their biomasses (see black arrows pointing down to such result for “Harvested Carnivores” in panels A and C of Fig. 4). The reverse occurred when increasing plankton subsidy, which dampened the effect of fishing on the biomass of harvested carnivores and filter-feeders. Interestingly, decreasing plankton subsidy also increased the biomass of harvested and non-harvested omnivores and herbivores (see results of subsection 2.3), which therefore dampened slightly the negative effect of fisheries on harvested omnivores and herbivores (Fig. 4 A and C).

Fisheries also affected the impacts of perturbing plankton subsidy on species biomasses. Increasing fishing increased the biomass of non-harvested species (see results of subsection 2.2) and, therefore, dampened the negative effects of altering plankton subsidy on the biomass of these species while intensifying the positive effects of altering plankton subsidy in those species (compare panels A and C with B and D of Fig. 4). Specifically, fisheries reversed the negative effect of increasing plankton subsidy on the biomass of non-harvested herbivores (see 4 black arrows pointing down to such result in panels A-C).

## DISCUSSION

An overall decrease in marine plankton productivity is expected over global scales^24,30,33^ as a result of climate change. Our results show that a decline in plankton productivity of the proportions expected with climate change can strongly impact an intertidal food-web in the Southern Pacific Coast. A decrease in plankton subsidy caused several species extinctions and shorten the food-web, with strong impacts on fisheries because of the biomass reduction of harvested species. Conversely, the simulated extinction of all harvested species caused null secondary extinctions, result that we also found in the subtidal food-web of the same Pacific Coast (see Supplementary Discussion and Supplementary Fig. S6). This despite artisanal fisheries harvesting on more than 20% of the food-web species (also highly connected species), which suggests that harvested species are embedded in redundant^51^ trophic interactions, conferring the food-web alternative routes of energy and stability^47,52^. These results, however, do not imply that local fisher communities will be similarly tolerant to the extinction of harvested species. The socio-economic system in which fishers are embedded will be directly impacted^53^ if resource management by local TURFs fails and drive the harvested species extinct (see Supplementary Discussion).

Our results also highlight the vulnerability of basal species to fishing. We found that basal species went extinct with an extraction above 0.3% of their available biomass. Harvested basal species are consumed by 2.5 more species than harvested consumers, and their intrinsic growth rate is 3 times lower than that of non-harvested basal species because they are the macroalgae that have the largest body size. Among the harvested basal species is kelp, which plays an important ecological and economical role. Kelp provides habitat structure and shelter to many species^12^ and via this non-trophic interaction, it promotes the biodiversity of coastal ecosystems^54^. Its commercial value is also high, with Chile being one of the main exploiters of kelp natural populations^55^. Kelp extraction in Chile is managed but hardly supervised^55^. Therefore, kelp’s high demand, high value, and low control, leave these algae prone to illegal fishing. In this context, we highlight the urgency of increasing supervision of kelp fisheries and enforcing their compliance with management plans.

We found that the effects on the food-web dynamics of coastal-productivity changes and artisanal fisheries interact, which reinforces the call made by previous studies^56–58^ that more research is needed to understand the interaction of several environmental stressors on ecosystems. Fisheries dampened the negative impacts of decreasing plankton productivity on non-harvested species, while plankton-productivity decline increased the sensitivity of harvested species to fishing. Previous work^59^ shows that human-gatherers enhance the species persistence of coastal marine ecosystems in the North Pacific. This suggests that, at least in the intertidal food-web studied here, small-scale artisanal fisheries play a similar role as human-gatherers in the North Pacific, that is, as keystone species to non-harvested species into the food-web.

Our study shows that the effects of climate change threaten the biodiversity of marine intertidal rocky-shore ecosystems as well as the services they provide, and emphasize the relevance of understanding and predicting the population dynamics of plankton and their impacts on entire food-webs (see Supplementary Discussion). New strategies for these new scenarios caused by climate change are needed to also protect the economy and wellbeing of the local communities dependent on these coastal ecosystems.

## METHODS

### Food-web description and the relative importance of harvested species to the food-web structure

We studied a well-resolved food-web of the intertidal rocky shore communities of the central coast of Chile^12^, which is harvested exclusively by small scale artisanal fisheries^11^. The web represents all species that are found to co-occur on wave exposed rocky platforms of central Chile, from the very low to the highest intertidal and is composed of 107 species (including a fisheries node), with 44% of its species corresponding to primary producers, 53% to invertebrates, and 3% to endotherm vertebrates. In the food-web, we consider as basal level all species of benthic primary producers plus plankton (phytoplankton + zooplankton, single node). Therefore, we represented filter-feeders (sessile filter-feeders + porcenallidae crabs) as specialist consumers of plankton and not as basal species (see detailed description of the food-web in Supplementary Material).

Species harvested by artisanal fisheries were identified using information from the Chilean national fishing service (www.sernapesca.cl) and previous work^13^. A high diversity of species distributed across all trophic levels are harvested by artisanal fisheries (red nodes in Fig. 1), including numerous species of macroalgae (n= 7), filter-feeders (n= 2), herbivores and omnivorous (n= 11), and carnivores (n= 2), totaling 22 species. Using the static approach (without population dynamics, see nest subsection), we compared the structure of the food-web with and without the harvested species to the distribution of 1000 food-web structures produced by randomly removing the same amount of harvested species (see more details about this method in supplementary materials).

### Static and dynamic approaches for evaluating food-web robustness

The static approach stems from network theory and analyzes the impacts of structural changes on food-webs represented by nodes (species) and links (interactions) that connect nodes, but ignores interaction strengths and population dynamics of interacting species^38^. In this approach, a non-basal species is considered extinct after a perturbation (defined here as a secondary extinction) if all its resource species (food) go extinct. Basal species are assumed to be autotrophs or otherwise obtain resources from outside the modeled web, e.g. through subsidies from other ecosystems and, therefore, do not experience extinctions unless directly removed (defined here as a primary extinctions). Thus, the static approach only considers extinctions produced by direct bottom-up effects. A dynamic approach considers not only the structure and intensity of interactions in a food-web, but also the changes in species abundances through time and the indirect and dynamic effects that these changes have on the abundances of other species^39–41,46,47^. A species is then considered to be secondarily extinct when its abundance drops below a threshold as a consequence of its population losses being higher than its population gains. Therefore, a dynamic approach can take into account both bottom-up and top-down effects of perturbations on the system, and both forces can contribute to produce secondary extinctions^39^. We use both the static network-based approach and a dynamic approach based on energy-transfer (see dynamic model below) to evaluate the impacts of artisanal fisheries and changes in primary productivity on the intertidal food-web.

### The dynamic model

The Allometric Trophic Network (ATN) model^49,50^ consists of two basic sets of equations, one set describing the biomass changes of primary producers (eq. 1) and the other describing that of consumers (eq. 2), where ***B*** is the biomass vector with the biomasses of every species population in the food-web and *B_i_* is the biomass of the population of species *i*, as follows:

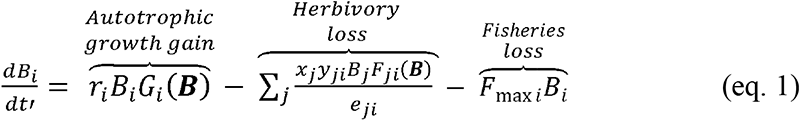

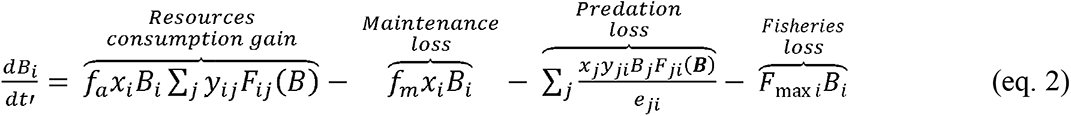

The biomass of producer *i* changes according to the balance of autotrophic growth gain and losses due to predation. The net autotrophic growth is determined by the logistic growth function *G_i_*(*B*) = 1 − (∑*_j=productores_c_ij_B_j_*)/*K*, where *r_i_* is the intrinsic growth rate of producer *i*, *c_ij_* is the inter-specific competition coefficient between producer *i* and *j*, and *K* is the total carrying capacity of primary producers in the system. The biomass loss of producer *i* by herbivory (caused by herbivores or omnivores) increases with the mass-specific metabolic (*x_j_*) and attack (*y_j_*) rates of consumer *i*, and decreases with the assimilation efficiency of consumer *i* for resource *j* (*e_ij_*). The consumers’ population dynamics (eq. 2) depend on their mass-specific metabolic rates (*x_j_*) and on the balance between biomass gains by resource consumption, biomass loss by metabolic maintenance, and biomass loss to predation. From the total amount of resources ingested by the consumer population *i*, ∑*_j_y_ij_F_ij_*(***B***), only a fraction *f_a_* is assimilated into consumer available energy for maintenance and biomass growth. The functional response *F_ij_*(***B***) determines the consumption rate of each consumer *i* for each resource *j*, defined by:

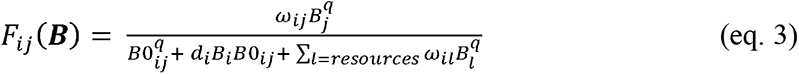

where *ꞷ_ij_* is the relative preference of consumer *i* for resource *j*, *q* controls the shape of eq. 3 which becomes an intermediate functional response between type II and type III when *q*=1.2 ^60^. *B*0_*ij*_ is the biomass of resource *j* at which the consumer *i* achieves half of its maximum consumption rate on resource *j*, and *d_i_* is the intra-specific interference of consumer *i* when it forages resource *j*. In Eq. 2, *f_m_* defines the fraction of the consumer biomass that is respired for maintenance of basal metabolism. *F_max_* defines the fraction of biomass *i* that is removed by small-scale artisanal fisheries. In the case of non-harvested species *F_max_* = 0.

The biological rates of production, R, metabolism, X, and maximum consumption, Y, follow a negative power law with the species body size, with an exponent -1.4[^61^]:

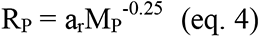

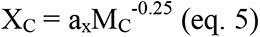

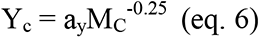

Where a_r_, a_x_, and a_y_ are allometric constants and the subscripts P and C denote producers and consumers, respectively. The timescale to examine the dynamics of the system is defined based on the primary producer with the highest mass-specific growth rate (reference species). The mass-specific growth rate and the metabolic rate of each species were normalized by the growth rate of the reference species, and the maximum consumption rate was normalized by each species’ metabolic rate^61^. These normalizations translate to the following expressions of intrinsic growth rate (*r_i_*), metabolic rate (*x_i_*), and maximum consumption rate (*y_i_*) of each species *i*:

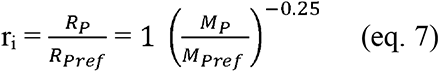

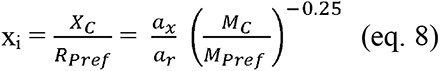

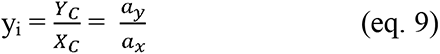

Since most benthic marine communities are characterized by the presence of sessile filter-feeders at the bottom, these communities are heavily ‘subsidized’ by the pelagic phytoplankton, which is captured by filter-feeders and transferred up the benthic food web^62^. In general, phytoplankton dynamics is thought to vary primarily due to ‘external processes’ (e.g. water advection, nutrient loadings, etc.,), including climate fluctuations^34^. To account for this phenomenon, our implementation of the ATN model assumed that the intertidal community is permanently subsidized by plankton biomass. Therefore, we modeled plankton dynamic as a primary producer (eq. 1) and following [75, 76] we incorporated a constant subsidy *s* into the plankton dynamics as:

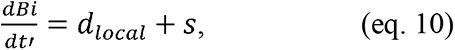

where *d_local_* represents plankton local dynamics (i.e., right hand of eq. 1).

To our knowledge, the ATN model we develop here is the largest empirical dynamic food-web model ever parameterized (our intertidal food-web contains 107 species, see model parametrization in supplementary materials and parameters values in Supplementary Table S4), which by itself represents an advancement in food web modeling. Moreover, the added realism of plankton subsidy allows us to simulate the effect of climate-change driven effects described above as the alteration of such subsidy.

### Food-web robustness to species extinctions

Using the static and dynamic approaches (see the model above), we evaluated the food-web robustness to species extinctions using four deletion sequences. First, we evaluated the food-web robustness to the extinction of harvested species by removing them in descending order of total catch amount (hereafter “harvesting” deletion sequence), according to the Chilean national fishing service (www.sernapesca.cl). Second, we performed three additional deletion sequences: (1) randomly (hereafter “random” deletion sequence), (2) from the most to the least connected species (hereafter “most-connected” deletion sequence^38,41^), and (3) from the most connected species that trophically support highly connected species to the least connected species supporting low connected species^42^. This last sequence causes the fastest route of collapse^42^ by first deleting the basal species that support most of the species in the food-web (hereafter “supporting-basal” deletion sequence). These last three deletion sequences allow us to compare the food-web sensitivity to the extinction of harvested species with that to the extinction of other species, and to identify the most important species for food-web persistence. For the harvesting deletion sequence, species were removed until all the harvested species were deleted, while for all other sequences the procedure was repeated until all species were removed (including the harvested species). In the case of species with an equal number of interactions, the removed species was chosen at random^40^.

To compare the food-web robustness across the different patterns of species deletion, except the “harvesting” deletion sequence, we use the R_50_ index^38^. For the harvesting deletion sequence, only the number of secondary extinctions was registered. The R_50_ index represents the proportion of species that have to be removed to cause the extinction of 50% of the species in the network (including primary and secondary extinctions). The highest and lowest possible values of R50 are 0.5 and 1/S, respectively (S is the number of species in the network), which are reached when no secondary extinctions are caused by species deletions and when only one primary species deletion is needed to cause the extinction of 50% of species, respectively. Thus, larger values of *R_50_* mean higher robustness. The static approach was simulated using the R package *NetworkExtinction*^63^, while the dynamical model was simulated using ODE45 in MATLAB.

For the dynamic approach, we first ran the dynamic model for 3650 time-steps which corresponds to 10 years, and ensures that the food-web reached a dynamic equilibrium. Then, we started the removal simulations. In each removal step, the number of extinct species was recorded after 10 years, when the system had reached, again, a steady state. A species was considered extinct if its biomass diminished to less than 10^-6^ [^64^]. Note that in all deletion sequences we removed the nodes from the food-web, so in the harvesting deletion sequence the F_max_ parameter in the ATN model is zero to all harvested species.

### Effects of artisanal fisheries on food-web dynamics

We assessed the effects of artisanal fisheries on food-web dynamics by simulating simultaneous fishing on all harvested species and assessing the subsequent effects on the biomass of all species in the food-web. We simulated three fishing scenarios, where we applied exploitation rates needed to decrease the biomass of all harvested species in 50%, 80%, and 100% (see F_max_ in Supplementary Table S3). Note that basal species required much lower exploitation rate to decrease their biomass (see discussion) than filter-feeders, herbivores, and other consumers, which means that harvested basal species were the most sensitive species to fishing. Note also that a biomass decrease of 100% does not necessarily mean that the harvested species go extinct, because the biomass available to be removed by fishing is the biomass that was produced a time step earlier (i.e., fishing exploitation is simulated as part of the population dynamics of harvested species, see eqs. 1 and 2). These three fishing scenarios allowed us to simulate an approximately well managed fisheries (which removes between 40 and 60% of biomass stock^5^), an overexploitation scenario (which removes 80%) and nearly extinction scenario, which allowed us to assess overall stability if all harvested species go extinct. For each fishing scenario, we first ran the model for 10 years (3650 time-steps) to ensure that the system reached a dynamic equilibrium. Then, we applied the biomass removal (*F*_maxi_*B*_i_ in eqs. 1 and 2) at each time step in the model to all harvested species simultaneously and we ran the food-web dynamics for another 3650 time-steps to reach post perturbation equilibrium, when final biomasses were considered “after perturbation” state.

In each treatment and for each species *i*, we evaluated the effect of the simulated scenario as:

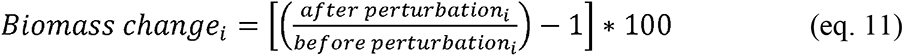

### Effects of plankton subsidy alteration on food-web dynamics

We assume that the food-web is subsidized by an external source, by including a subsidy in the plankton node, which is considered to be controlled by advective processes, unaffected by local benthic consumption. This represents well the situation of most marine benthic ecosystems^62^. To compare the top-down effects of fisheries against bottom-up variation in productivity, we perturbed the plankton subsidy to simulate climate-induced changes in plankton productivity. We simulated both a decrease and an increase in plankton subsidy, as both long-term increased and decreased productivity has been documented to occur in the Humboldt Ecosystem^34^. We used three different perturbation intensities, decreasing or increasing basal subsidy in 50%, 80% and 100%. Note that, a 100% in the basal plankton subsidy decreasing does not translate into plankton extinction (Fig. 3C). A variation of 50% of the basal subsidy is in the order of natural seasonal variability of net primary productivity in central Chile^31^. We assumed that a variation above 50% simulates the effects of extreme changes of plankton subsidy due to climate change, and also, such magnitudes allow comparable perturbation intensities to those used to assess the impacts of fisheries on the food-web dynamics (see previous section). In each scenario, we first ran the model for 3650 time-steps to ensure that the system reached a dynamic equilibrium, and the final species biomasses obtained were considered “*before perturbation*” state. Then, we reduced/increased plankton subsidy at each time step and ran the model for another 3650 time-steps to reach post perturbation equilibrium. The final biomasses were considered “*after perturbation*” state. Changes in biomass were expressed as shown in Eq.11.

### Interacting effects of fisheries and plankton-subsidy alteration on food-web dynamics

To evaluate combined effects of fisheries and climate-induced changes in plankton subsidy, we simulated both fishing on all harvested species and simultaneously altered plankton subsidy. We used the three fisheries scenarios (i.e., 50%, 80% and 100% biomass removed) and crossed these scenarios with each of the six productivity scenarios (i.e., increasing or decreasing plankton subsidy in 50%, 80% and 100%). In each treatment, we first ran the model for 3650 time-steps and the final species biomasses obtained were considered “*before perturbation*” state. Then, we applied a given plankton subsidy scenario, and at the same time, we started the fishing simulations. Changes in biomass were expressed as shown in Eq.11.

## DATA AVAILABILITY

Simulation code and the Chilean intertidal data will be available upon acceptance at the repository https://github.com/fsvaldovinos/Chilean_Fisheries. The Chilean intertidal food-web and species body sizes can also be found in^12^

## Acknowledgments

We thank Kayla R.S. Hale, Paul Glaum, and Valentin Coco for technical help on modelling and MATLAB programming, as well as, Jonathan Morris, Joseph Hartert, and Feng-Hsun Chang for insightful discussions (all members of Valdovinos lab). We thank Mercedes Pascual and Stefano Allesina for valuable comments and for sharing with us their code of their eigenvector-based algorithm. We also thank E.A. Wieters for relevant opinions about empirical data. MA was funded by CONICYT doctoral fellowship 21160860. PM acknowledges support from project AFB-17008.

## Author Contributions

M.I.A., S.A.N., P.A.M., and F.S.V. conceived the study. M.I.A. and F.S.V. formulated the dynamic model, designed and implemented simulations, analyzed results, and wrote the first draft of the manuscript. S.A.N. E.A.W and A.P. created the trophic interaction database and led food web compilation. M.I.A. and D.C. created the code to perform the static extinction analysis. All authors contributed to the final version of the paper.

## Additional Information

Competing financial interests: The authors declare no competing financial interests.

## References

1. Barnosky, A. D. et al. Has the Earth’s sixth mass extinction already arrived?. Nature 471, 51–57 (2011).

2. McCauley, D. J. et al. Marine defaunation: Animal loss in the global ocean. Science 347, 1255641–1255641 (2015).

3. Chapin III, F. S. et al. Consequences of changing biodiversity. Nature 405, 234–242 (2000).

4. Díaz, S., Fargione, J., Chapin, F. S. & Tilman, D. Biodiversity Loss Threatens Human Well-Being. PLoS Biol. 4, e277 (2006).

5. Worm, B. et al. Rebuilding Global Fisheries. Science 325, 578–585 (2009).

6. Defeo, O. & Castilla, J. C. More than One Bag for the World Fishery Crisis and Keys for Co-management Successes in Selected Artisanal Latin American Shellfisheries. Rev. Fish Biol. Fish. 15, 265–283 (2005).

7. Pauly, D. & Zeller, D. Catch reconstructions reveal that global marine fisheries catches are higher than reported and declining. Nat. Commun. 7, 10244 (2016).

8. Defeo, O. et al. Co-management in Latin American small-scale shellfisheries: assessment from long-term case studies. Fish Fish. 17, 176–192 (2016).

9. Gelcich, S. et al. Fishers’ perceptions on the Chilean coastal TURF system after two decades: problems, benefits, and emerging needs. Bull. Mar. Sci. 93, 53–67 (2017).

10. Castilla, J. C., Gelcich, S. & Defeo, O. Successes, Lessons, and Projections from Experience in Marine Benthic Invertebrate Artisanal Fisheries in Chile. in Fisheries Management (eds. McClanahan, T. R. & Castilla, J. C.) 23–42 (Blackwell Publishing Ltd, 2007). doi:10.1002/9780470996072.ch2.

11. Gelcich, S. et al. Navigating transformations in governance of Chilean marine coastal resources. Proc. Natl. Acad. Sci. 107, 16794–16799 (2010).

12. Kéfi, S. et al. Network structure beyond food webs: mapping non-trophic and trophic interactions on Chilean rocky shores. Ecology 96, 291–303 (2015).

13. Pérez-Matus, A. et al. Temperate rocky subtidal reef community reveals human impacts across the entire food web. Mar. Ecol. Prog. Ser. 567, 1–16 (2017).

14. Pérez-Matus, A., Carrasco, S. A., Gelcich, S., Fernandez, M. & Wieters, E. A. Exploring the effects of fishing pressure and upwelling intensity over subtidal kelp forest communities in Central Chile. Ecosphere 8, e01808 (2017).

15. Gelcich, S. et al. Territorial User Rights for Fisheries as Ancillary Instruments for Marine Coastal Conservation in Chile: Gelcich et al. Conserv. Biol. 26, 1005–1015 (2012).

16. Oyanedel, R., Keim, A., Castilla, J. C. & Gelcich, S. Illegal fishing and territorial user rights in Chile: Illegal Fishing. Conserv. Biol. 32, 619–627 (2018).

17. Donlan, C. J., Wilcox, C., Luque, G. M. & Gelcich, S. Estimating illegal fishing from enforcement officers. Sci. Rep. 10, 12478 (2020).

18. Andreu-Cazenave, M., Subida, M. D. & Fernandez, M. Exploitation rates of two benthic resources across management regimes in central Chile: Evidence of illegal fishing in artisanal fisheries operating in open access areas. PLOS ONE 12, e0180012 (2017).

19. Castilla, J. C. Coastal marine communities: trends and perspectives from human-exclusion experiments. Trends Ecol. Evol. 14, 280–283 (1999).

20. Somero, G. N. The physiology of climate change: how potentials for acclimatization and genetic adaptation will determine ‘winners’ and ‘losers’. J. Exp. Biol. 213, 912–920 (2010).

21. Hoegh-Guldberg, O. & Bruno, J. F. The Impact of Climate Change on the World’s Marine Ecosystems. Science 328, 1523–1528 (2010).

22. Brose, U. et al. Climate change in size-structured ecosystems. Philos. Trans. R. Soc. B Biol. Sci. 367, 2903–2912 (2012).

23. Doney, S. C. et al. Climate Change Impacts on Marine Ecosystems. Annu. Rev. Mar. Sci. 4, 11–37 (2012).

24. Kwiatkowski, L., Aumont, O. & Bopp, L. Consistent trophic amplification of marine biomass declines under climate change. Glob. Change Biol. 25, 218–229 (2019).

25. Bakun, A. Coastal Ocean Upwelling. Science 247, 198–201 (1990).

26. Bakun, A., Field, D. B., Redondo-Rodriguez, A. & Weeks, S. J. Greenhouse gas, upwelling-favorable winds, and the future of coastal ocean upwelling ecosystems. Glob. Change Biol. 16, 1213–1228 (2010).

27. Thiel, M. et al. The Humboldt current system of northern and central Chile: oceanographic processes, ecological interactions and socioeconomic feedback. in Oceanography and Marine Biology (eds. Gibson, R., Atkinson, R. & Gordon, J.) vol. 20074975 195–344 (CRC Press, 2007).

28. Morales, C., Hormazabal, S., Andrade, I. & Correa-Ramirez, M. Time-Space Variability of Chlorophyll-a and Associated Physical Variables within the Region off Central-Southern Chile. Remote Sens. 5, 5550–5571 (2013).

29. Aiken, C. M., Navarrete, S. A. & Pelegrí, J. L. Potential changes in larval dispersal and alongshore connectivity on the central Chilean coast due to an altered wind climate. J. Geophys. Res. 116, G04026 (2011).

30. Blanchard, J. L. et al. Potential consequences of climate change for primary production and fish production in large marine ecosystems. Philos. Trans. R. Soc. B Biol. Sci. 367, 2979–2989 (2012).

31. Testa, G., Masotti, I. & Farías, L. Temporal variability in net primary production in an upwelling area off central Chile (36°S). Front. Mar. Sci. 5, 179 (2018).

32. Batten, S. D. et al. A Global Plankton Diversity Monitoring Program. Front. Mar. Sci. 6, 321 (2019).

33. Chust, G. et al. Biomass changes and trophic amplification of plankton in a warmer ocean. Glob. Change Biol. 20, 2124–2139 (2014).

34. Weidberg, N. et al. Spatial shifts in productivity of the coastal ocean over the past two decades induced by migration of the Pacific Anticyclone and Bakun’s effect in the Humboldt Upwelling Ecosystem. Glob. Planet. Change 193, 103259 (2020).

35. Aguirre, C., García-Loyola, S., Testa, G., Silva, D. & Farias, L. Insight into anthropogenic forcing on coastal upwelling off south-central Chile. Elem Sci Anth 6, 59 (2018).

36. Valdovinos, F. S. Mutualistic networks: moving closer to a predictive theory. Ecol. Lett. 22, 1517–1534 (2019).

37. Pascual, M. & Dunne, J. A. Ecological Networks: Linking Structure to Dynamics in Food Webs (Santa Fe Institute Studies on the Sciences of Complexity). (Oxford University Pres, 2006).

38. Dunne, J. A., Williams, R. J. & Martinez, N. D. Network structure and biodiversity loss in food webs: robustness increases with connectance. Ecol. Lett. 5, 558–567 (2002).

39. Curtsdotter, A. et al. Robustness to secondary extinctions: Comparing trait-based sequential deletions in static and dynamic food webs. Basic Appl. Ecol. 12, 571–580 (2011).

40. Ramos-Jiliberto, R., Valdovinos, F. S., Moisset de Espanés, P. & Flores, J. D. Topological plasticity increases robustness of mutualistic networks: Interaction rewiring in mutualistic networks. J. Anim. Ecol. 81, 896–904 (2012).

41. Valdovinos, F. S., Moisset de Espanés, P., Flores, J. D. & Ramos-Jiliberto, R. Adaptive foraging allows the maintenance of biodiversity of pollination networks. Oikos 122, 907–917 (2013).

42. Allesina, S. & Pascual, M. Googling Food Webs: Can an Eigenvector Measure Species’ Importance for Coextinctions?. PLoS Comput. Biol. 5, e1000494 (2009).

43. de Santana, C., Rozenfeld, A., Marquet, P. & Duarte, C. Topological properties of polar food webs. Mar. Ecol. Prog. Ser. 474, 15–26 (2013).

44. Albert, R., Jeong, H. & Barabási, A. Error and attack tolerance of complex networks. Nature 406, 378–382 (2000).

45. Ives, A. R. & Cardinale, B. J. Food-web interactions govern the resistance of communities after non-random extinctions. Nature 429, 174–177 (2004).

46. Rebolledo, R., Navarrete, S. A., Kéfi, S., Rojas, S. & Marquet, P. A. An open-system approach to complex biological networks. SIAM J. Appl. Math. 79, 619–640 (2019).

47. McCann, K. S. The diversity–stability debate. Nature 405, 228–233 (2000).

48. Williams, Rich J. Network 3D: visualizing and modelling food webs and other complex networks. Microsoft Res. Camb. UK (2010).

49. Richard, J. W., Brose, U. & Martinez, N. D. Homage to Yodzis and Innes 1992: Scaling up feeding-based population dynamics to complex ecological networks. in From Energetics to Ecosystems: The Dynamics and Structure of Ecological Systems 37–51 (Springer Netherlands, 2006). doi:10.1007/978-1-4020-5337-5_2.

50. Boit, A., Martinez, N. D., Williams, R. J. & Gaedke, U. Mechanistic theory and modelling of complex food-web dynamics in Lake Constance: Mechanistic modelling of complex food web dynamics. Ecol. Lett. 15, 594–602 (2012).

51. Jordán, F., Okey, T. A., Bauer, B. & Libralato, S. Identifying important species: Linking structure and function in ecological networks. Ecol. Model. 216, 75–80 (2008).

52. Arim, M. & Marquet, P. A. Intraguild predation: a widespread interaction related to species biology: Intraguild predation. Ecol. Lett. 7, 557–564 (2004).

53. Castilla, J. C. & Fernandez, M. Small-scale benthic fisheries in Chile: on co-management and sustainable use of benthic invertebrates. Ecol. Appl. 8, S124–S132 (1998).

54. Teagle, H., Hawkins, S. J., Moore, P. J. & Smale, D. A. The role of kelp species as biogenic habitat formers in coastal marine ecosystems. J. Exp. Mar. Biol. Ecol. 492, 81–98 (2017).

55. Vásquez, J. A. The Brown Seaweeds Fishery in Chile. in Fisheries and Aquaculture in the Modern World (ed. Mikkola, H.) (InTech, 2016). doi:10.5772/62876.

56. He, Q. & Silliman, B. R. Climate Change, Human Impacts, and Coastal Ecosystems in the Anthropocene. Curr. Biol. 29, R1021–R1035 (2019).

57. Brown, C. J., Saunders, M. I., Possingham, H. P. & Richardson, A. J. Interactions between global and local stressors of ecosystems determine management effectiveness in cumulative impact mapping. Divers. Distrib. 20, 538–546 (2014).

58. Crain, C. M., Kroeker, K. & Halpern, B. S. Interactive and cumulative effects of multiple human stressors in marine systems. Ecol. Lett. 11, 1304–1315 (2008).

59. Dunne, J. A. et al. The roles and impacts of human hunter-gatherers in North Pacific marine food webs. Sci. Rep. 6, 21179 (2016).

60. Williams, R. J. Effects of network and dynamical model structure on species persistence in large model food webs. Theor. Ecol. 1, 141–151 (2008).

61. Brose, U., Williams, R. J. & Martinez, N. D. Allometric scaling enhances stability in complex food webs. Ecol. Lett. 9, 1228–1236 (2006).

62. Menge, B. A. & Menge, D. N. L. Dynamics of coastal meta-ecosystems: the intermittent upwelling hypothesis and a test in rocky intertidal regions. Ecol. Monogr. 83, 283–310 (2013).

63. Ávila-Thieme, M. I., Corcoran, D., Valdovinos, F. S., Navarrete, S. A. & Marquet, P. A. NetworkExtinction: Extinction Simulation in Food Webs. (R package version 0.1.3., 2018).

64. Schneider, F. D., Brose, U., Rall, B. C. & Guill, C. Animal diversity and ecosystem functioning in dynamic food webs. Nat. Commun. 7, 12718 (2016).

