## Supplementary material for "Alteration of coastal productivity and artisanal fisheries interact to affect a marine food-web"

### **Food-web description**

We studied the food-web of the intertidal rocky-shore ecosystem of central Chile. This food-web is influenced by the highly productive Humboldt Current System^1^ (HCS) and harvested exclusively by small scale artisanal fisheries^2^. The web represents all species that are found to co-occur on wave exposed rocky platforms of central Chile, from the very low to the highest intertidal and is composed of 107 species (including fisheries node), with 44% of its species corresponding to primary producers, 53% to invertebrates, and 3% to endotherm vertebrates^3^. A total of 1381 consumption type interactions have been documented in the web, with approximately linear increase in species degree (number of interactions per node) and species rank^3^, and connectance of 0.12. In the food-web, we consider as basal level all species of benthic primary producers (e.g. algae) plus plankton (phytoplankton + zooplankton, single node). Therefore, we represented filter-feeders (sessile filter-feeders + porcenallidae crabs) as specialist consumers of plankton and not as a basal species, which could be alternative approach if they were not connected with the plankton. Benthic diatoms were considered as an independent node, separate from plankton. Detailed methods and general description of food web attributes, as well as patterns of spatial variability can be found in [3–5].

### **Relative importance of harvested species for the food-web structure**

Using the static approach (without population dynamics), we compared the structure of the food-web with and without the harvested species to the distribution of 1000 food-web structures produced by randomly removing the same amount of harvested species. This comparison used: number of species (S), number of trophic links (L), connectance (L/S^2^), number of omnivores, and averaged trophic level (MeanSWTL). We calculated all these structural metrics using cheddar package in R. After removing all harvested species using our static approach, the food-web resulted in lower link density (*L*/*S*, where *L* and *S* are numbers of trophic interactions and species, respectively) and connectance (C = *L*/*S* ^2^ ) than removing the same number of species at random (Table S2). This decrease in connectance is counter-intuitive because the pure removal of species should increase connectance based on its mathematical definition. Therefore, this result shows that disproportionately more interactions than species are lost with the removal of harvested species. In addition, the removal of harvested species decreased the fraction of omnivore species and the mean trophic level but this decrease is not different to that produced by the random removal of the same number of species (Table S2), suggesting that the impact of artisanal fisheries may not be strong enough to destabilize^6^ and shorten^7^ the intertidal food-web. Finally, the number of secondary extinctions produced by the removal of harvested species was less than the average number of secondary extinctions obtained when removing the same number of species but selected at random (Table S2). These results reinforce the observation that artisanal fisheries share prey with other consumers that also forage on other non-harvested species in the food-web, so alternative resources remain when all harvested species go extinct.

### **Model Parametrization**

Supplementary Table S4 shows all the model parameters with their initial values and descriptions. We used empirical measures of species body size, which was extracted from [3], to allometrically parameterize intrinsic growth rate of autotroph, as well as metabolic rate, and maximum consumption rate of each species (see Methods in main text).

The initial biomass was estimated from density (mobile species + cnidaria) and the surface cover (sessile species) recorded during six years of sampling in the Chilean marine reserve “Estacion Costera de Investigaciones Marinas” (ECIM) at the central-south of Chile. The density of mobile species was multiplied by the species average body size. For sessile species, the surface cover was multiplied by the weight of each species per unit of cover (e.g. [8]).

We calculated the community-level carrying capacity, *K,* by dividing all the primary producers into six functional groups (microalgae, ephemerals, corticates, crust, corallines, and kelps)^4^. In each functional group, we multiplied the biomass of the species exhibiting the highest growth rate with the number of species that compose its functional group. Finally, we summed the resulting biomass over all functional groups.

Based on empirical experiments in aquatic^9^ and terrestrial ecosystems^10^, we assumed that the half-saturation density parameter (B0) decreases while the trophic levels increase in 10^3^ order of magnitude. We use the values of Boit et al.^11^ as a reference value. To herbivores, we fit the B0 to ensure their persistence. For the other parameters, we used values from Calbet, A. & Saiz ^9^ (see Supplementary Table S4).

### **Supplementary Discussion**

Fisheries are commonly associated with negative impacts on ecosystems, especially on the food-web structure, total biomass, and species recovery^6,7,12^. The present study shows that these effects are not necessarily general at a food-web scale. We found that in a rocky-shore intertidal food-web, the simulated extinction of all harvested species caused null secondary extinctions. This despite artisanal fisheries harvesting on more than 20% of the food-web species, which are also highly connected species. In addition, we found that this food-web was highly vulnerable to the decrease of plankton productivity, which is one of the outcomes expected to happen as consequence of climate change ^13–16^. Finally, we found that artisanal fisheries might contribute to dampening the negative consequences of plankton-productivity decrease by increasing the biomass of non-harvested species. In the following paragraphs, we expand on these results and contextualize them with prior literature.

The impact on the food-web structure of all harvested species going extinct (i.e., food-web shortened and connectance decreased, Supplementary Table S2) caused null secondary extinctions. This suggests that harvested species are embedded in redundant^17^ trophic interactions. In addition, the high food-web robustness suggests that the exploitative competition between artisanal fisheries with harvested and non-harvested species for common resources (Fig. 1C) might be weak, as consumers have wider diets that can buffer the loss of harvested species. The high redundancy of trophic interactions is explained by its high levels of omnivory^18^, generalist consumers^19^, and a high proportion of transient and weak links^20^. These attributes confer food-webs alternative routes of energy and stability^21,22^. These results, however, do not imply that local fisher communities will be similarly tolerant to the extinction of harvested species or to future perturbations. The socio-economic system in which fishers are embedded will be directly impacted^23^ if resource management by local TURFs fails and drive the harvested species extinct. Fishers would need to harvest on new species as alternative resources to maintain their livelihood. In our analysis, the loss of harvested species caused the loss of an important amount of redundant links (Table S2), which suggests that the resulting food-web contains a greater predominance of functional than redundant links, and therefore, less robust to further species extinctions^24^. Therefore, fishers exploiting new species will impact the food-web in ways we did not explore here.

In the rocky intertidal ecosystems we studied in this work, artisanal fishermen obtain their resources through hand-picking and use them for self-subsistence^25^. Therefore, resource availability in intertidal ecosystems plays an important role for the poorest fishermen. Artisanal fishers with more means apply most of their fishing gears (e.g., diving, spearing, and pot trapping) in subtidal-shallow ecosystems^25^, from where they harvest ~20% of species in the food-web (Supplementary Fig. S6A). For the above, we repeated our static extinction analysis in the shallow-subtidal marine food-web and found that, as in the intertidal food-web, the subtidal food-web is robust to the loss of all harvested species (Supplementary Fig. S6B). This suggests that similar mechanisms explaining the high food-web robustness of the intertidal ecosystem against the extinction of harvested species, explain the high food-web robustness of the subtidal. Moreover, as in the intertidal food-web, we found that plankton was the most important group (node) for the subtidal food-web persistence. Analyzing the effect of artisanal fisheries on the subtidal food-web with a dynamic approach seems an important next step to understand how anthropogenic activities as well as bottom-up and top-down forces affect coastal food-webs.

A concerning effect of climate change is the alteration of plankton productivity. This can be caused by the physicochemical changes in coastal waters triggered by warmer waters^13^ and by an intensification of upwelling-favorable winds^26,27^, accompanied with a decrease (or increase) of nutrients given by an intensification of the warm (or cold) phase of ENSO^28–30^. The importance of plankton is well-known as the energy supply of food-webs, as well as essential for sustaining fisheries^31^. We found that plankton is the most important food-web component for species persistence. Plankton is consumed by filter-feeders and any alteration of plankton subsidy affects the biomass of all the species in the food-web. On the one hand, a decrease in plankton subsidy caused the intertidal food-web to shorten, with strong impacts on fisheries because of the biomass reduction of harvested species. Similar results were found when climate change effects were simulated as an increase of biological rates^32,33^ caused by temperature raises, which suggests that our results will be magnified if we were to consider the alteration of biological rates. In addition, a reduction in plankton productivity may reduce the recruitment of species (as plankton composition also include larvae of several species^34^), which might cause more secondary extinctions than we found here. On the other hand, an increase in plankton subsidy negatively impacted the biomass of a higher number of species but in smaller magnitude than the decrease of plankton subsidy. Moreover, an enrichment of nutrients can increase the arrival of new species^35^ or the recurrence of harmful algal blooms with a devastating effect on local food-webs^36^. Thus, if we consider these factors, we would expect an intensification in the negative consequences observed in this study.

Limitations of our research mostly consist of factors not included in our modeling approach. For example, artisanal fisheries harvest kelp which plays an important non-trophic role by providing habitat structure and shelter to many species^3^. Other factors not considered in this study are local spatial features such as the enclosed bay with internal circulation and larval retention^37^ as well as upwelling zones^38^, which can affect species recruitment and affect our results. Similarly, temporal variability and other stochasticity sources associated with global change (e.g., invasive species, pathogens spread, habitat degradation, and several climatic stressors^39,40^) can also change the relative importance of species in food-webs, making an open-system approach^41^ to ecological networks an important next step in the area. For example, the extinction of all harvested species might release several ecological niches and, consequently, fisheries might increase the species invasion. Moreover, as invasive species are characterized by generalist foraging habits and lacking predators^42^, we might find negative consequences in the abundance of local non-harvested species. Thus, our results should be interpreted with care, and the positive effects of fisheries do not mean that fisheries can indiscriminately exploit these ecosystems. These effects might depend on adaptive prey-switching behavior (mechanism not considered here), allowing them to use alternative or new resources in response to changes in abundances of other species in the community, and with that rewire food-web^43^ and stabilize populations dynamics^44^.

### **References***:*

1.	Thiel, M. *et al.* The Humboldt current system of northern and central Chile: oceanographic processes, ecological interactions and socioeconomic feedback. in *Oceanography and Marine Biology* (eds. Gibson, R., Atkinson, R. & Gordon, J.) vol. 20074975 195–344 (CRC Press, 2007).

2.	Gelcich, S. *et al.* Navigating transformations in governance of Chilean marine coastal resources. *Proceedings of the National Academy of Sciences* **107**, 16794–16799 (2010).

3.	Kéfi, S. *et al.* Network structure beyond food webs: mapping non-trophic and trophic interactions on Chilean rocky shores. *Ecology* **96**, 291–303 (2015).

4.	Kéfi, S., Miele, V., Wieters, E. A., Navarrete, S. A. & Berlow, E. L. How Structured Is the Entangled Bank? The Surprisingly Simple Organization of Multiplex Ecological Networks Leads to Increased Persistence and Resilience. *PLoS Biol* **14**, e1002527 (2016).

5.	Lurgi, M. *et al.* Geographical variation of multiplex ecological networks in marine intertidal communities. *Ecology* (2020) doi:10.1002/ecy.3165.

6.	Kuparinen, A., Boit, A., Valdovinos, F. S., Lassaux, H. & Martinez, N. D. Fishing-induced life-history changes degrade and destabilize harvested ecosystems. *Sci Rep* **6**, 22245 (2016).

7.	Pauly, D. Fishing Down Marine Food Webs. *Science* **279**, 860–863 (1998).

8.	Wieters, E. A., Broitman, B. R. & Brancha, G. M. Benthic community structure and spatiotemporal thermal regimes in two upwelling ecosystems: Comparisons between South Africa and Chile. *Limnol. Oceanogr.* **54**, 1060–1072 (2009).

9.	Calbet, A. & Saiz, E. Effects of trophic cascades in dilution grazing experiments: from artificial saturated feeding responses to positive slopes. *Journal of Plankton Research* **35**, 1183–1191 (2013).

10.	Mulder, C. & Hendriks, A. J. Half-saturation constants in functional responses. *Global Ecology and Conservation* **2**, 161–169 (2014).

11.	Boit, A., Martinez, N. D., Williams, R. J. & Gaedke, U. Mechanistic theory and modelling of complex food-web dynamics in Lake Constance: Mechanistic modelling of complex food web dynamics. *Ecology Letters* **15**, 594–602 (2012).

12.	Jackson, J. B. C. Historical Overfishing and the Recent Collapse of Coastal Ecosystems. *Science* **293**, 629–637 (2001).

13.	Kwiatkowski, L., Aumont, O. & Bopp, L. Consistent trophic amplification of marine biomass declines under climate change. *Glob Change Biol* **25**, 218–229 (2019).

14.	Blanchard, J. L. *et al.* Potential consequences of climate change for primary production and fish production in large marine ecosystems. *Phil. Trans. R. Soc. B* **367**, 2979–2989 (2012).

15.	Weidberg, N. *et al.* Spatial shifts in productivity of the coastal ocean over the past two decades induced by migration of the Pacific Anticyclone and Bakun’s effect in the Humboldt Upwelling Ecosystem. *Global and Planetary Change* **193**, 103259 (2020).

16.	Chust, G. *et al.* Biomass changes and trophic amplification of plankton in a warmer ocean. *Glob Change Biol* **20**, 2124–2139 (2014).

17.	Jordán, F., Okey, T. A., Bauer, B. & Libralato, S. Identifying important species: Linking structure and function in ecological networks. *Ecological Modelling* **216**, 75–80 (2008).

18.	Pérez-Matus, A., Carrasco, S. A., Gelcich, S., Fernandez, M. & Wieters, E. A. Exploring the effects of fishing pressure and upwelling intensity over subtidal kelp forest communities in Central Chile. *Ecosphere* **8**, e01808 (2017).

19.	Camus, P. A., Arancibia, P. A. & Ávila-Thieme, M. I. A trophic characterization of intertidal consumers on Chilean rocky shores. *Rev. biol. mar. oceanogr.* **48**, 431–450 (2013).

20.	Lopez, D. N., Camus, P. A., Valdivia, N. & Estay, S. A. High temporal variability in the occurrence of consumer-resource interactions in ecological networks. *Oikos* **126**, 1699–1707 (2017).

21.	McCann, K. S. The diversity–stability debate. *Nature* **405**, 228–233 (2000).

22.	Arim, M. & Marquet, P. A. Intraguild predation: a widespread interaction related to species biology: Intraguild predation. *Ecology Letters* **7**, 557–564 (2004).

23.	Castilla, J. C. & Fernandez, M. Small-scale benthic fisheries in Chile: on co-management and sustainable use of benthic invertebrates. *Ecological Applications* **8**, S124–S132 (1998).

24.	Allesina, S., Bodini, A. & Pascual, M. Functional links and robustness in food webs. *Phil. Trans. R. Soc. B* **364**, 1701–1709 (2009).

25.	Defeo, O. & Castilla, J. C. More than One Bag for the World Fishery Crisis and Keys for Co-management Successes in Selected Artisanal Latin American Shellfisheries. *Rev Fish Biol Fisheries* **15**, 265–283 (2005).

26.	Aguirre, C., García-Loyola, S., Testa, G., Silva, D. & Farias, L. Insight into anthropogenic forcing on coastal upwelling off south-central Chile. *Elem Sci Anth* **6**, 59 (2018).

27.	Belmadani, A., Echevin, V., Codron, F., Takahashi, K. & Junquas, C. What dynamics drive future wind scenarios for coastal upwelling off Peru and Chile?. *Clim Dyn* **43**, 1893–1914 (2014).

28.	Wang, Y., Luo, Y., Lu, J. & Liu, F. Changes in ENSO amplitude under climate warming and cooling. *Clim Dyn* **52**, 1871–1882 (2019).

29.	Cai, W. *et al.* Increased variability of eastern Pacific El Niño under greenhouse warming. *Nature* **564**, 201–206 (2018).

30.	Cai, W. *et al.* Increased frequency of extreme La Niña events under greenhouse warming. *Nature Clim Change* **5**, 132–137 (2015).

31.	Batten, S. D. *et al.* A Global Plankton Diversity Monitoring Program. *Front. Mar. Sci.* **6**, 321 (2019).

32.	Brose, U. *et al.* Climate change in size-structured ecosystems. *Phil. Trans. R. Soc. B* **367**, 2903–2912 (2012).

33.	Fussmann, K. E., Schwarzmüller, F., Brose, U., Jousset, A. & Rall, B. C. Ecological stability in response to warming. *Nature Clim Change* **4**, 206–210 (2014).

34.	Hays, G., Richardson, A. & Robinson, C. Climate change and marine plankton. *Trends in Ecology & Evolution* **20**, 337–344 (2005).

35.	Jochum, M., Schneider, F. D., Crowe, T. P., Brose, U. & O’Gorman, E. J. Climate-induced changes in bottom-up and top-down processes independently alter a marine ecosystem. *Phil. Trans. R. Soc. B* **367**, 2962–2970 (2012).

36.	Hallegraeff, G. M. A review of harmful algal blooms and their apparent global increase. *Phycologia* **32**, 79–99 (1993).

37.	Morgan, S. G., Fisher, J. L., Miller, S. H., McAfee, S. T. & Largier, J. L. Nearshore larval retention in a region of strong upwelling and recruitment limitation. *Ecology* **90**, 3489–3502 (2009).

38.	Ospina-Alvarez, A., Weidberg, N., Aiken, C. M. & Navarrete, S. A. Larval transport in the upwelling ecosystem of central Chile: The effects of vertical migration, developmental time and coastal topography on recruitment. *Progress in Oceanography* **168**, 82–99 (2018).

39.	Barnosky, A. D. *et al.* Has the Earth’s sixth mass extinction already arrived?. *Nature* **471**, 51–57 (2011).

40.	McCauley, D. J. *et al.* Marine defaunation: Animal loss in the global ocean. *Science* **347**, 1255641–1255641 (2015).

41.	Rebolledo, R., Navarrete, S. A., Kéfi, S., Rojas, S. & Marquet, P. A. An open-system approach to complex biological networks. *SIAM J. Appl. Math.* **79**, 619–640 (2019).

42.	Sakai, A. K. *et al.* The population biology of invasive species. *Annu. Rev. Ecol. Syst.* **32**, 305–332 (2001).

43.	Thierry, A. *et al.* Adaptive foraging and the rewiring of size-structured food webs following extinctions. *Basic and Applied Ecology* **12**, 562–570 (2011).

44.	Valdovinos, F. S., Ramos-Jiliberto, R., Garay-Narváez, L., Urbani, P. & Dunne, J. A. Consequences of adaptive behaviour for the structure and dynamics of food webs: adaptive behaviour in food webs. *Ecology Letters* **13**, 1546–1559 (2010).

45.	Gómez-Canchong, P., Quiñones, R. A. & Brose, U. Robustness of size–structure across ecological networks in pelagic systems. *Theor Ecol* **6**, 45–56 (2013).

46.	Brose, U., Williams, R. J. & Martinez, N. D. Allometric scaling enhances stability in complex food webs. *Ecol Letters* **9**, 1228–1236 (2006).

47.	Testa, G., Masotti, I. & Farías, L. Temporal variability in net primary production in an upwelling area off central Chile (36°S). *Front. Mar. Sci.* **5**, 179 (2018).

48.	Williams, Rich J. Network 3D: visualizing and modelling food webs and other complex networks. *Microsoft Research, Cambridge, UK* (2010).

### **Tables**

**Table S1**. Rank of the 30 most connected species of the intertidal food-web

| **Rank** | **Species name** | **Harvested** | **Total number of interactions (degree)** |
| --- | --- | --- | --- |
| 1 | *Fissurella limbata* | yes | 67 |
| 2 | *Fissurella crassa* | yes | 63 |
| 3 | *Acanthopleura echinata* | yes | 60 |
| 4 | *Fissurella costata* | yes | 60 |
| 5 | *Chiton granosus* | yes | 59 |
| 6 | *Chiton latus* | no | 57 |
| 7 | *Chiton cummingii* | no | 56 |
| 8 | *Enoplochiton niger* | no | 56 |
| 9 | *Chaetopleura peruviana* | no | 52 |
| 10 | *Fissurella cummingii* | yes | 47 |
| 11 | *Heliaster helianthus* | no | 43 |
| 12 | *Scurria araucana* | no | 41 |
| 13 | *Siphonaria lesoni* | no | 41 |
| 14 | Gulls | no | 41 |
| 15 | *Acanthocyclus gayi* | no | 40 |
| 16 | *Fissurella maxima* | yes | 39 |
| 17 | *Tegula atra* | yes | 39 |
| 18 | *Tonicia benaventii* | no | 39 |
| 19 | *Tonicia chilensis* | no | 39 |
| 20 | *Tonicia elegans* | no | 39 |
| 21 | *Scurria ceciliana* | no | 38 |
| 22 | *Scurria variabilis* | no | 38 |
| 23 | *Fissurella picta* | yes | 37 |
| 24 | *Fissurella puhlcra* | yes | 37 |
| 25 | *Scurria plana* | no | 37 |
| 26 | *Scurria viridula* | no | 37 |
| 27 | *Acanthocyclus hassleri* | no | 36 |
| 28 | *Balanus laevis* | no | 36 |
| 29 | *Nothobalanus flosculus* | no | 36 |
| 30 | *Lottia orbigny* | no | 35 |

**Table S2.** Structural properties of the intertidal food-web before and after removing all 22 harvested species (i.e., “After

non-random removal”), and after removing the same number of species randomly (i.e., “After random removal”). The “After random removal” column shows the mean and 95% confidence interval of structural property values of 1000 iterations of randomly removing 22 species out of the 107 species in the food-web. The fourth column represents the percentage of change of each structural property, calculated as [(“after non-random removal” – “before any species removal”)/ “before any species removal”]. The sixth column indicate whether the change of the structural properties produced by the loss of harvested species (non-random deletion) is different to the changes produced by random deletions. SWTLMean is the short-weighted trophic level of a food-web.

| Structural property | Before any species removal | After  non-random removal | Percentage of change (%) | After  random  removal | Non-random  vs  random removal |
| --- | --- | --- | --- | --- | --- |
| Richness | 107 | 85 | -21 | 82 ± 0.4 | ≠ |
| Number of links | 1381 | 718 | -48 | 819 ± 6 | ≠ |
| Connectance | 0.12 | 0.10 | -18 | 0.13 ± 0 | ≠ |
| %-Omnivore | 37 | 32 | -12 | 32 ± 0 | = |
| MeanSWTL | 1.64 | 1.62 | -1 | 1.62 ± 0.01 | = |

**Table S3.** F_max_ values used to produce the total biomass decrease of the harvested basal species, harvested filter-feeders, and the other harvested species in a 50%, 80% and 100%. F_max_ = 0.001 means that fisheries remove 0.1% of the available biomass of the harvested species; while F_max_ = 1 means that fisheries remove 100% of the available biomass of the harvested species.

| Type of harvested species | Biomass decrease of interest in harvested species | F_max_ value that produce the biomass decrease of interest |
| --- | --- | --- |
| Basal species |  |  |
|  | 50% | 0.00125 |
|  | 80% | 0.0022 |
|  | 100% | 0.01 |
| Filter-feeders and herbivores |  |  |
|  | 50% | 0.23 |
|  | 80% | 0.8 |
|  | 100% | 1 |
| Others consumers |  |  |
|  | 50% | 0.23 |
|  | 80% | 0.8 |
|  | 100% | 1 |

**Table S4**. Initial parameter values used in our version of the Allometric Trophic Network (ATN) model. In references column ECIM refers to the coastal marine research station of the Pontificia Universidad Catolica de Chile.

| Parameter | Unit of measurement | Definition | Initial values  min., max | References |
| --- | --- | --- | --- | --- |
| B | g / m^2^ | Population abundance | 1.26 x 10^-4^, 112107 | Empirical values from ECIM, Diatoms, and Plankton values from [^45^] |
| M | g | Body mass | 1 x 10^-5^, 500 | [3] |
| r | 1 / day | Mass-specific growth rate of basal species | 0.1075, 3.76 | Calculated using [^46^] |
| x | 1 / day | Mass-specific metabolic rate of consumers | 0.7284, 70.96 | Calculated using [^46^] |
| y | - | Maximum consumption rate | 1, 5.8 | Calculated using [^46^] |
| K | g / m^2^ | Carrying capacity of basal species | 176299 | K adapted from [11] |
| c | - | Competition coefficient of basal species | 1 | [11] |
| fa | - | The fraction of biomass that is assimilated from consumer | 0.4 | [11] |
| fm | - | The fraction of biomass that is respired to metabolic maintenance | 0.1 | [11] |
| e | - | Assimilation efficiency | 0.45, 0.85 | [^46^] |
| d | m^2^ / g | Intra-specific interference | 0.5 | [11] |
| q | - | Functional response | 1.2 | [11] |
| w | - | Resources preference | 1/n_resources_ | [11] |
| p | - | Fractions of shared resources | 0, 1 | [11] |
| B0 | g / m^2^ | Half-saturation density | 150, 15000 | Adapted from [11] |
| s | g / m^2^ | The subsidy that is considered in plankton dynamic. | 12% of the initial biomass | [^47^] |

**Table S5**. Species name corresponding to the species number shown in the x-axis of Supplementary Figs. S2 and S4.

| Code | Specie Name | Code | Specie Name | Code | Specie Name |
| --- | --- | --- | --- | --- | --- |
| 1 | *Acanthina monodon* | 47 | *Scurria viridula* | 93 | *Peysonella* spp. |
| 2 | *Concholepas concholepas* | 48 | *Scurria zebrina* | 94 | *Plocamium cartilageneum* |
| 3 | *Acanthopleura echinata* | 49 | *Siphonaria lessoni* | 95 | *Prionitis* spp. |
| 4 | *Chiton granosus* | 50 | *Echinolittorina peruviana* | 96 | *Gastroclonium cyclindricum* |
| 5 | *Fissurella costata* | 51 | *Austrolittorina araucana* | 97 | *Rhodymenia* sp. |
| 6 | *Fissurella crassa* | 52 | *Onchidella* sp*.* | 98 | *Schottera nicaensis* |
| 7 | *Fissurella cummingi* | 53 | *Balanus laevis* | 99 | *Schyzimenia doryophora* |
| 8 | *Fissurella limbata* | 54 | *Jhelius cirratus* | 100 | *Trematocarpus* spp. |
| 9 | *Fissurella maxima* | 55 | *Nothobalanus flosculus* | 101 | *Corallina offcinalis v*ar. Chilensis |
| 10 | *Fissurella picta* | 56 | *Nothochthamalus scabrosus* | 102 | *Hildenbrandia lecanelieri* |
| 11 | *Fissurella puhlcra* | 57 | *Brachidontes granulata* | 103 | *Lithothamnion* spp*.* |
| 12 | *Scurria scurra* | 58 | *Perumytilus purpuratus* | 104 | *Ralfsia californica* |
| 13 | *Tegula atra* | 59 | *Semimytilus algosus* | 105 | Benthic diatoms |
| 14 | *Austromegabalanus psittacus* | 60 | *Allelopetrolisthes punctatus* | 106 | Plankton |
| 15 | *Pyura chilensis* | 61 | *Petrolisthes spinifrons* |  |  |
| 16 | *Durvillaea antarctica* | 62 | *Petrolisthes angulosus* |  |  |
| 17 | *Lessonia nigrescens* | 63 | *Petrolisthes tuberculatus* |  |  |
| 18 | *Gelidium rex* | 64 | *Petrolisthes tuberculosus* |  |  |
| 19 | *Sarcothalia* spp. | 65 | *Phragmatopoma* spp*.* |  |  |
| 20 | *Mazzaella laminarioides* | 66 | *Bryopsis* spp. |  |  |
| 21 | *Pyropia* spp. | 67 | *Centroceras* spp. |  |  |
| 22 | *Ulva rigida* | 68 | *Ceramium* spp*.* |  |  |
| 23 | Gulls | 69 | *Chaetomorpha* spp. |  |  |
| 24 | *Cinclodes nigrofumosus* | 70 | *Cladophora* spp*.* |  |  |
| 25 | *Anthotoe* spp. | 71 | *Ectocarpus silicosus* |  |  |
| 26 | *Bunodactis* spp*.* | 72 | *Enteromorpha compressa* |  |  |
| 27 | *Oulactis concinnata* | 73 | *Halopteris funicularis* |  |  |
| 28 | *Parantheopsis* spp. | 74 | *Polysiphonia* spp. |  |  |
| 29 | *Phymactis* spp. | 75 | *Rhizoclonium* ambiguum |  |  |
| 30 | *Trimusculus peruvianus* | 76 | *Scythosiphon lomentaria* |  |  |
| 31 | *Heliaster helianthus* | 77 | *Ulvella* spp. |  |  |
| 32 | *Stichaster striatus* | 78 | *Adenocystis utricularis* |  |  |
| 33 | *Acanthocyclus gayi* | 79 | *Ahnfeltiopsis* spp*.* |  |  |
| 34 | *Acanthocyclus hassleri* | 80 | *Chondrus canaliculatus* |  |  |
| 35 | *Chaetopleura peruviana* | 81 | *Codium dimorpha* |  |  |
| 36 | *Chiton cummingsi* | 82 | *Colpomenia phaeodactyla* |  |  |
| 37 | *Chiton latus* | 83 | *Colpomenia sinuosa* |  |  |
| 38 | *Enoplochiton niger* | 84 | *Gelidium* spp*.* |  |  |
| 39 | *Tonicia lineolata* | 85 | *Glossophora kunthii* |  |  |
| 40 | *Tonicia chilensis* | 86 | *Grateloupia* spp. |  |  |
| 41 | *Tonicia elegans* | 87 | *Gymnogongrus furcellatus* |  |  |
| 42 | *Lottia orbignyi* | 88 | *Laurencia chilensis* |  |  |
| 43 | *Scurria araucana* | 89 | *Montemaria horridula* |  |  |
| 44 | *Scurria ceciliana* | 90 | *Nothogenia* spp. |  |  |
| 45 | *Scurria plana* | 91 | *Petalonia Fascia* |  |  |
| 46 | *Scurria variabilis* | 92 | *Petroglossum* spp*.* |  |  |

**Fig. S1**.

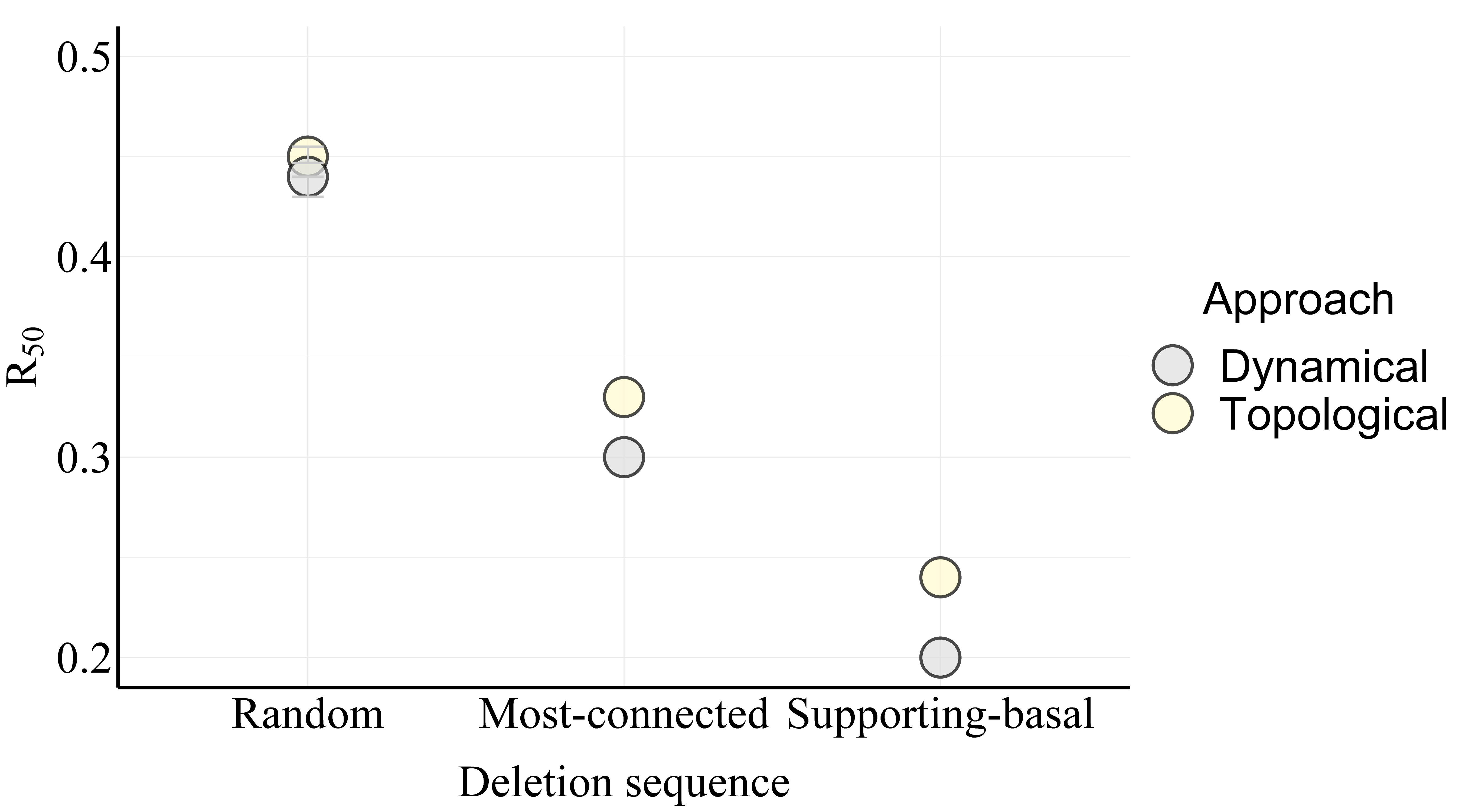

**Figure S1**. Food-web robustness (R_50_) to three deletion sequences and using a static (yellow circle) vs. dynamical (grey circle) approach. The performed deletion sequences removed species: (1) randomly (hereafter “random”), (2) from the most to the least connected species (hereafter “most-connected”), and (3) from the most connected species that trophically support highly connected species to the least connected species supporting low connected species (hereafter “Supporting-basal”). In the case of the random deletion sequence, the circles represent the average and the error bars the ± 95. C.I. of 1000 random deletion sequences.

**Fig. S2**

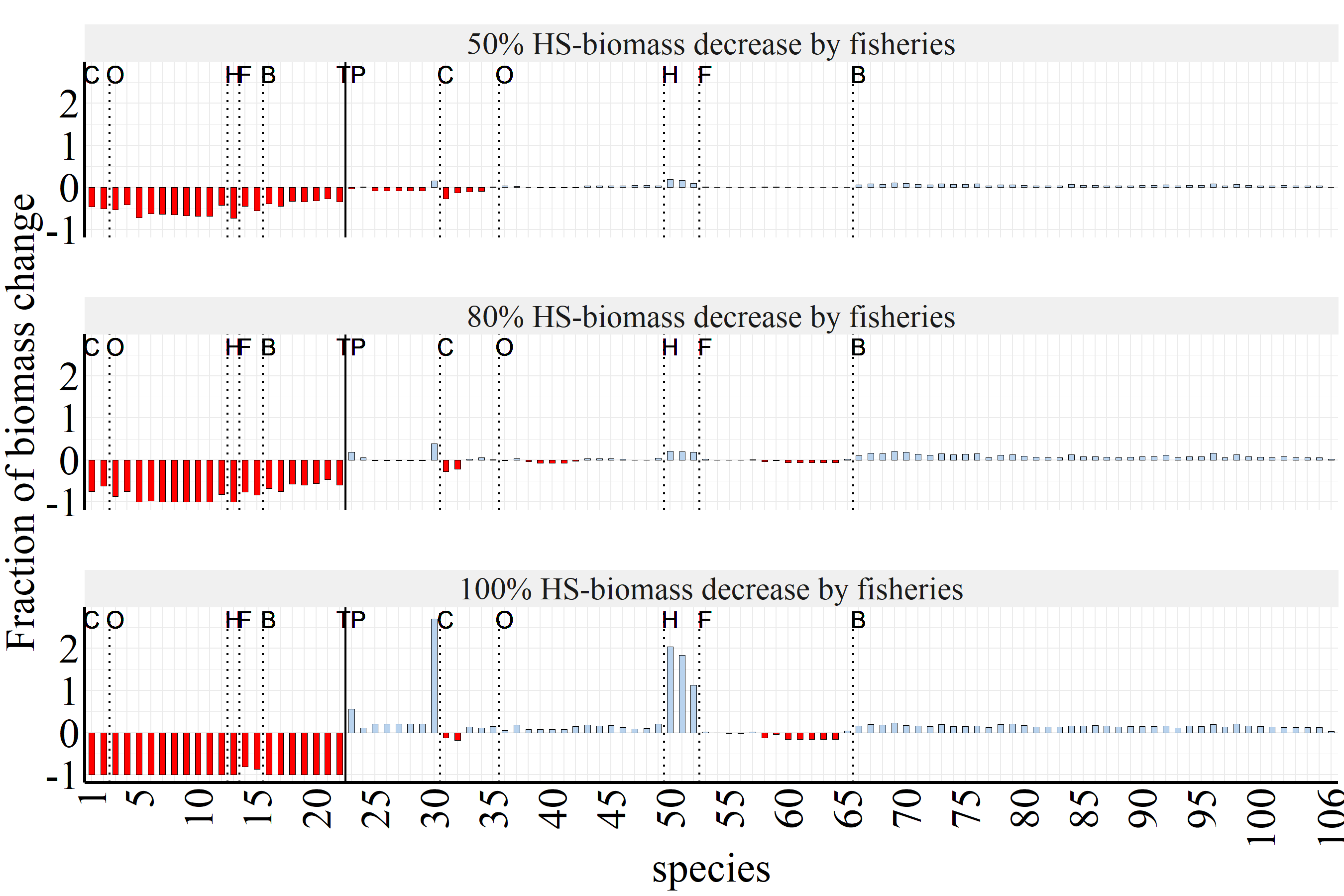

(A)

(B)

(C)

**Figure S2.** Fraction of biomass change (y-axis) of each species (x-axis) of the intertidal food-web after simultaneously reducing the biomass of all harvested species in a 50% (A), 80% (B) and 100% (C). Red bars represent negative effects on species biomass, while blue bars represent positive effects. From the bold vertical line to the left, the figure shows all the harvested species (HS). From the bold line to the right, the figure shows all the non-harvested species. Species are organized by trophic level (TP: top-predators, C: carnivores, O: omnivores, H: herbivores, F: filter-feeders, B: basal species) and their identity can be found by matching their number id with the numbers in Supplementary Table S5.

**Fig. S3**

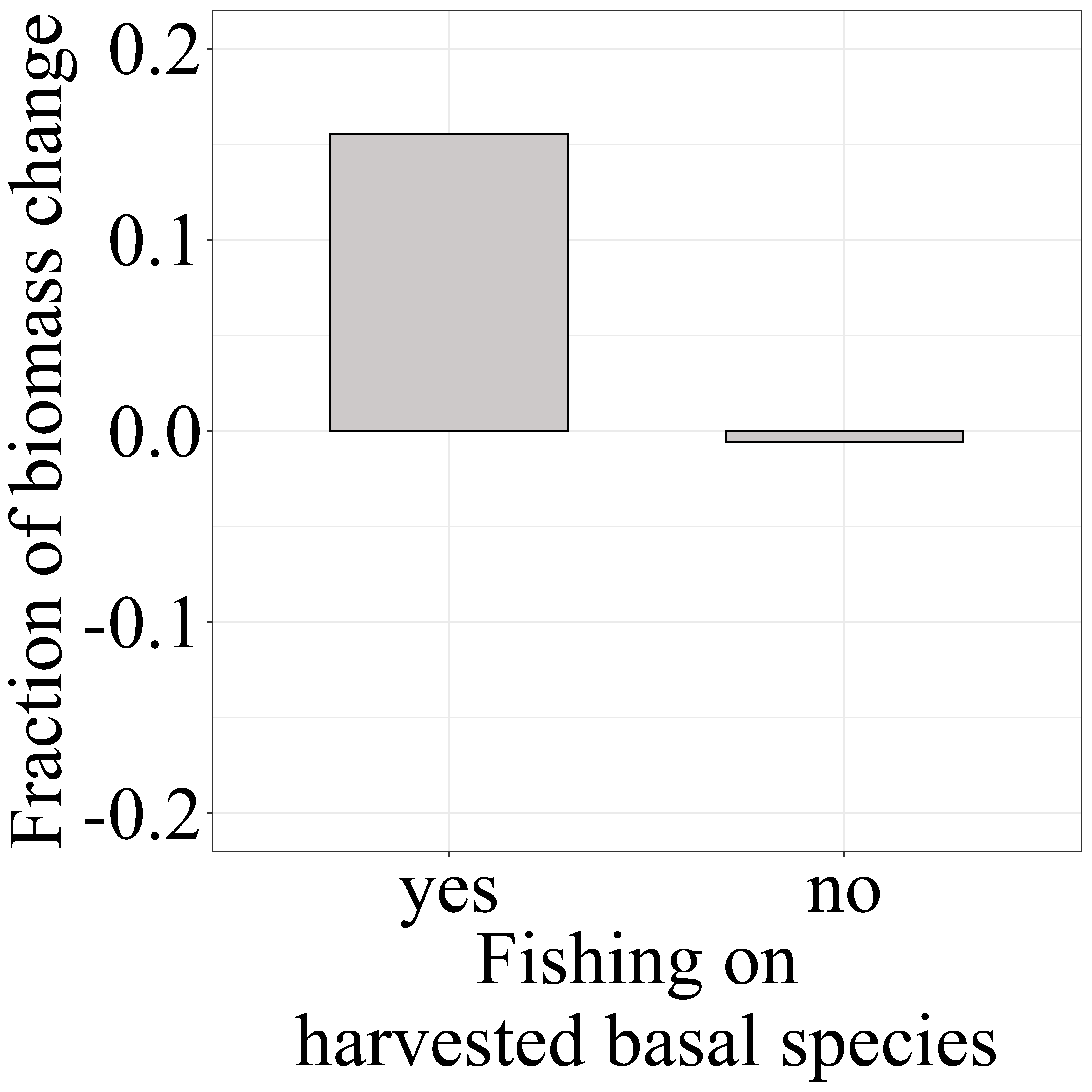

**Figure S3.** Contribution of harvested basal species over the biomass of non-harvested basal species. The figure represents the fraction of the total biomass change of non-harvested basal species (y-axis) after fishing with the maximum exploitation rate (F_max_ = 1) for all harvested species, including (yes) and without including (no) harvested macroalgae in the list of harvested species.

**Fig. S4**

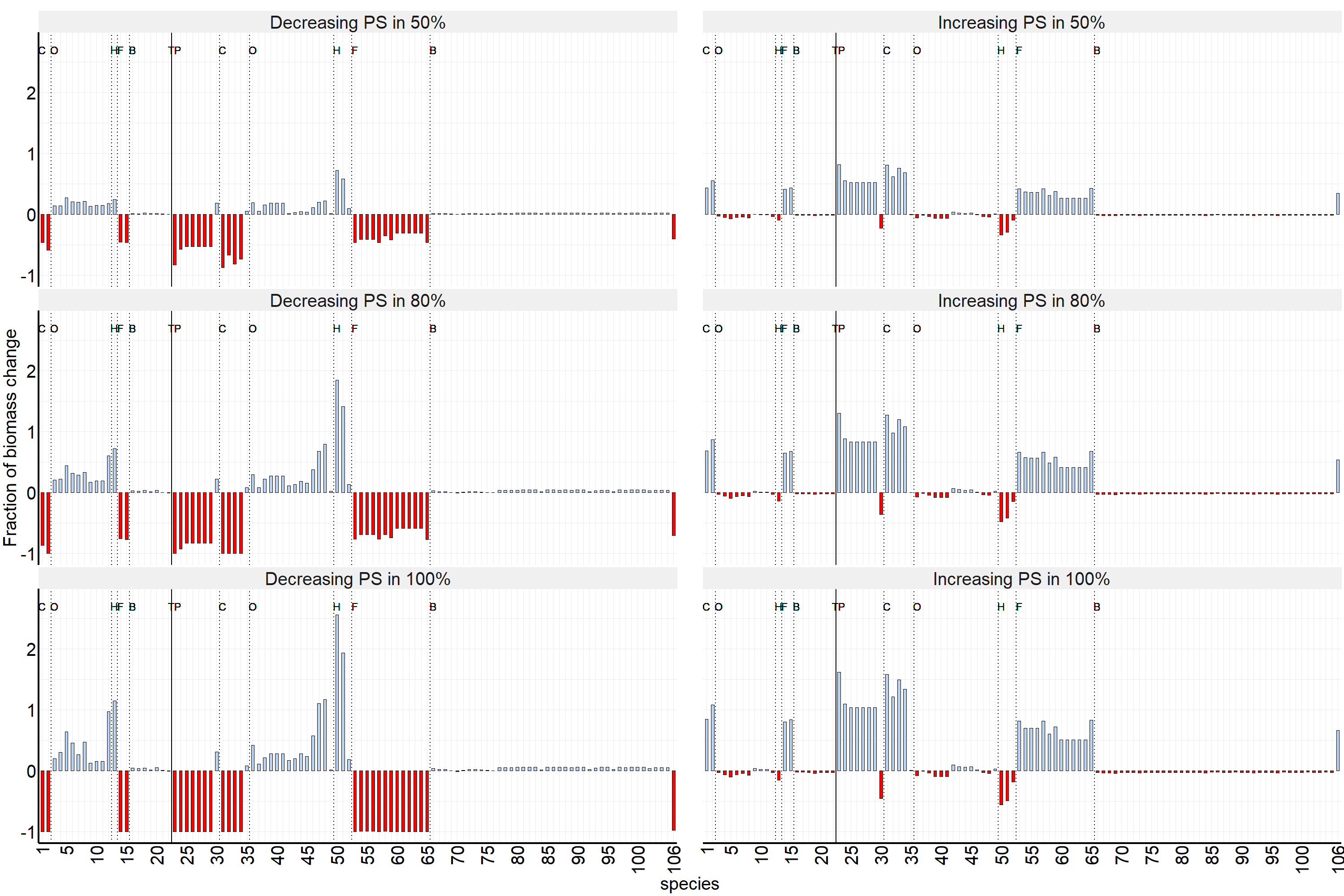

(A)

(B)

(C)

(D)

(E)

(F)

**Figure S4.** Fraction of biomass change (y-axis) of each species (x-axis) of the intertidal food-web after decreasing (A,C,E) and increasing (B,D,F) plankton subsidy (species 106) in 50% (A,B), 80% (C,D), 100% (E,F) of their basal productivity. The red bar represents negative effects on species biomass, while blue bars represent positive effects. From the bold vertical line to the left, the figure shows all the harvested species. From the bold line to the right, the figure shows all the non-harvested species. Species are organized by trophic level and their identity can be found by matching their number id with the numbers in Supplementary Table S5.

**Fig. S5**

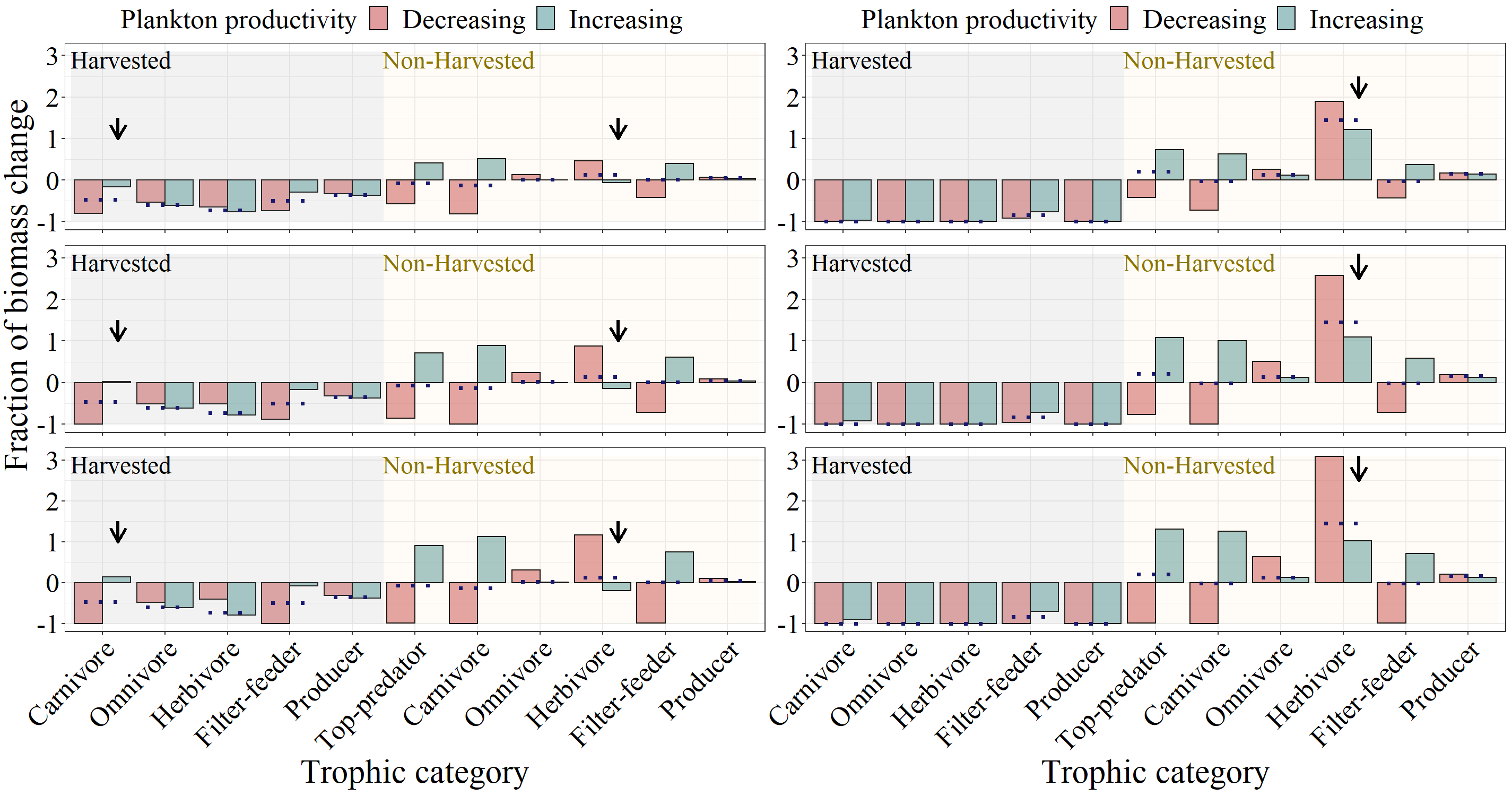

50% of PP

100% of PP

80% of PP

**(E)**

**(F)**

**(C)**

**(A)**

100% HS-biomass decrease by fisheries

50% HS-biomass decrease by fisheries

**(D)**

**(B)**

**Figure S5.** Combined effects of artisanal fisheries and plankton-productivity alterations on food-web dynamics. Fraction of total biomass change (y-axis) of each trophic category (x-axis) after decreasing (red bars) and increasing (blue bars) the plankton productivity (PP) in 50% (A and B), 80% (C and D), and 100% (E and F), and after decreasing the biomass of all harvested species (HS) in a 50% (A and C) and in a 100% (B and D). The grey and yellow shading represent the biomass change of harvested and non-harvested species, respectively. The arrows highlight the most remarkable changes between the two levels of plankton subsidy perturbation and the two levels of fishing. The dotted lines represent the independent effect fishing (i.e., without plankton subsidy perturbation) on the biomass of each trophic category as a reference point.

**Fig. S**
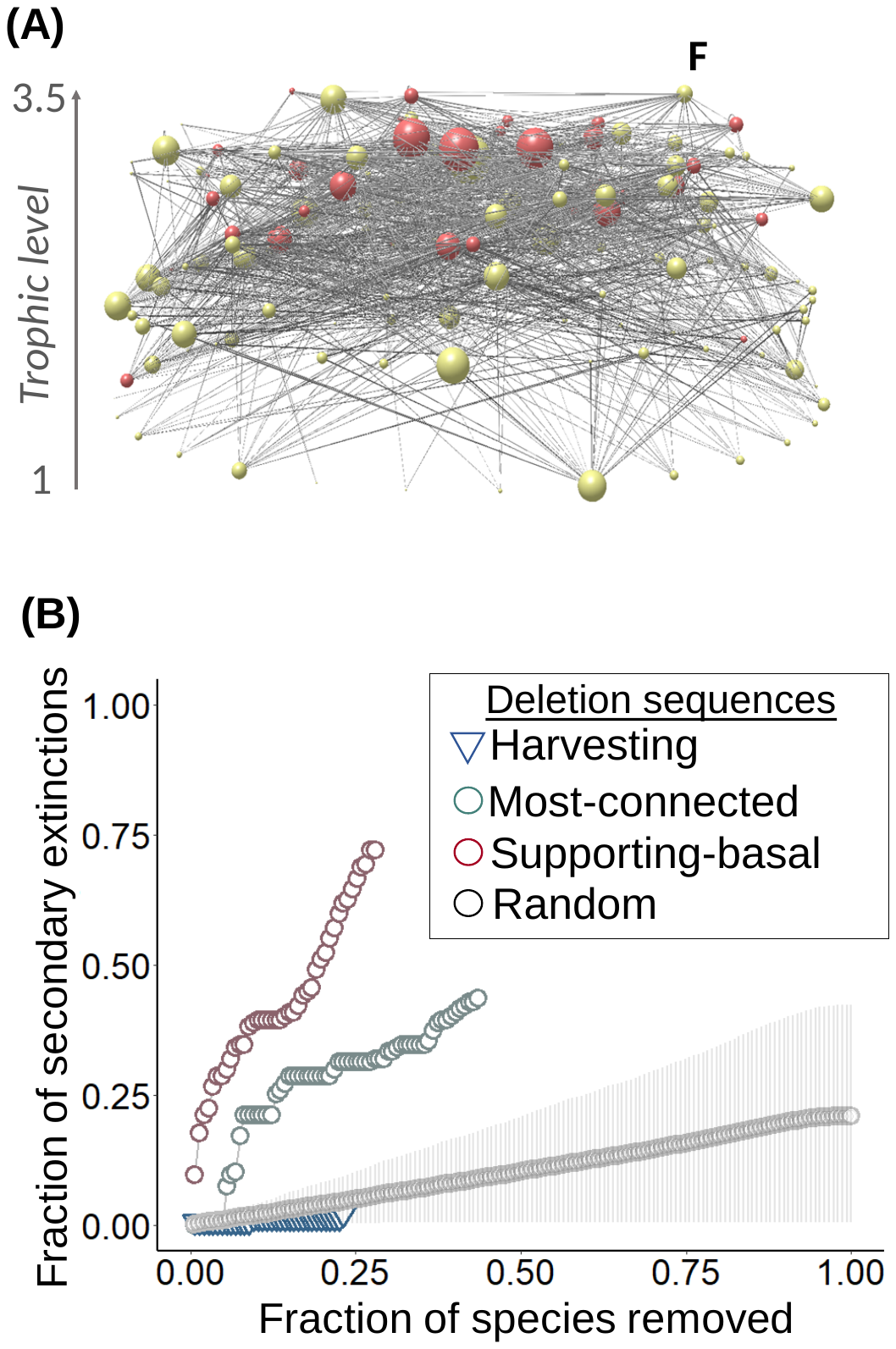
**6**

**Figure S6.** Shallow-subtidal marine Chilean food-web. (A) Color node represents the harvested (red) and non-harvested (yellow) species. Letter F represents the fisheries node. Node size represents the number of trophic interactions (degree) of each node. Bottom nodes represent basal species, and the nodes of the top represent top predators. The left axes represent the trophic level (SWTL) from the minimum to the maximum trophic level. Drawn using Network3D software^48^. Harvested species was recognized from the official governmental webpage of the Chilean fishing service ([www.sernapesca.cl](http://www.sernapesca.cl)). (B) Fraction of secondary extinctions (y-axis) produced in this food-web after the sequential removal of species (x-axis) with a static approach. Gray and red circle represent most-connected and supporting-basal deletion sequence, while blue tringle represents harvesting deletion sequence. In the random deletion sequence, circles represent the average and the error bars represent the 95% confidence interval of 1000 simulations.
